# Technical and biological sources of unreliability of Infinium probes on Illumina Methylation microarrays

**DOI:** 10.1101/2023.03.14.532595

**Authors:** Tatiana Nazarenko, Charlotte D. Vavourakis, Allison Jones, Iona Evans, Lena Schreiberhuber, Christine Kastner, Isma Ishaq-Parveen, Elisa Redl, Antony W. Watson, Kirsten Brandt, Clive Carter, Alexey Zaikin, Chiara Herzog, Martin Widschwendter

## Abstract

The Illumina Methylation array platform has facilitated countless epigenetic studies on DNA methylation (DNAme) in health and disease, yet relatively few studies have so studied its reliability, i.e., the consistency of repeated measures. Here we focus on the reliability of both type I and type II Infinium probes. We propose a method for excluding unreliable probes based on dynamic thresholds for mean intensity (MI) and ‘unreliability’, estimated by probe-level simulation of the influence of technical noise on methylation β-values using the background intensities of negative control probes. We validate our method in several datasets, including Illumina MethylationEPIC BeadChip v1.0 data from paired whole blood samples taken six weeks apart. Our analysis revealed that specifically probes with low MI exhibit higher β-value variability between repeated samples. MI was associated with the number of C-bases in the respective probe sequence and correlated negatively with unreliability scores. The unreliability scores were substantiated through validation in a new EPIC v1.0 (blood and cervix) and a publicly available 450k (blood) dataset, as they effectively captured the variability observed in β-values between technical replicates. Finally, despite promising higher robustness, the newer version v2.0 of the MethylationEPIC BeadChip retained a substantial number of probes with poor unreliability scores. To enhance current pre-processing pipelines, we developed an R package to calculate MI and unreliability scores and provide guidance on establishing optimal dynamic score thresholds for a given data set.

## INTRODUCTION

DNA methylation (DNAme) is a chemical modification of DNA that entails the addition of a methyl group to a cytosine (C) residue resulting in 5-methylcytosine, and most commonly occurs in the context of CpG dinucleotides in humans (1). The study of epigenetics and DNAme has become one of the most topical areas of genomic research in recent years, both from a functional point of view and a clinical perspective, owing to its potential application in cancer risk prediction and early detection strategies (2, 3).

The two most widely used techniques to study epigenome-wide DNAme are whole-genome bisulphite sequencing (WGBS) and Illumina methylation arrays. Both technologies require bisulfite pre-treatment of DNA to enable distinction of methylated from unmethylated cytosine residues in the context of CpG dinucleotides. Whereas WGBS provides information regarding the DNAme status of a series of linked CpGs, the Illumina methylation arrays allow a more affordable and high-throughput assessment of the methylation status of a subset single CpGs dinucleotides throughout the genome.

The Illumina BeadArray technology has undergone substantial re-development over the years and the total number of CpGs that can be simultaneously analysed has increased substantially from ∼25,000 in 2008 (HumanMethylation27KBeadChip), to ∼485,000 in 2011 (HumanMethylation450K BeadChip), to over ∼850,000 CpG sites in 2016 (MethylationEPIC BeadChip v1.0), and finally to over ∼935,000 CpG sites in 2022 when the MethylationEPIC BeadChip v2.0 was released. Illumina Methylation microarrays include two different types of bead chemistry, Infinium type I and II probes (4, 5). Type I probes have two separate probe sequences per CpG dinucleotide (one each for methylated and unmethylated CpGs), whereas type II probes have just one probe sequence per CpG dinucleotide. Consequently, type II probes take up much less physical space on the arrays than type I probes and are the most abundant type on the latest Illumina EPIC arrays, constituting ∼85% of all probes. For type II probes, discrimination of methylated (M) versus unmethylated (U) alleles is made possible by single nucleotide primer extension which results in the incorporation of Cy3 or Cy5 labelled nucleotides into the target sequence and emitting green or red fluorescence, respectively. For type I probes, discrimination of methylated versus unmethylated alleles is made by constructing corresponding probes sequences (M and U) which are measured in the same channel, either Red or Green. Further we will distinguish between them as type I-Red and I-Green probes respectively. The level of methylation at specific CpG sites is expressed as *Beta* (β-value), which represents a constant from ‘0’ (unmethylated) to ‘1’ (fully methylated) and can be written as:

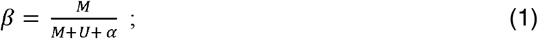

with α a small positive constant (typically 100) added to the equation to avoid dividing by zero when both M and U signals are equal to 0. If β = 0, then the interrogated CpG is unmethylated (there is no M signal), if β = 1, then the interrogated CpG is methylated (there is no U signal).

When assessing Illumina Methylation array data, basic pre-processing steps would typically include identifying probes and/or samples with a low signal to noise intensity which should be excluded (6), correcting for background intensity and dye bias (7), performing within-array normalization to reduce differences in beta-distributions obtained from Infinium I and II probes (8), and imputing missing data (9). For this, several established methods have been benchmarked and implemented into pre-processing pipelines available as R packages, such as *minfi* (10), *ChAMP* (11) and the latest *ENmix* (12). Additionally, previous studies have identified the necessity to exclude low-specificity probes that can bind to multiple sequences within the genome, as well as probes that contain genetic variants in their underlying sequence (5, 13). Lastly, a recent study by Sugden *et al*. (14) identified a large set of ‘unreliable’ probes that poorly reproduced methylation values when samples from the same DNA source were measured either on the HumanMethylation450K or MethylationEPIC BeadChip. However, to date comprehensive understanding of the factors which influence the reliability of Illumina array probes is lacking (where *reliability* refers to the ability to reproduce data). This has substantial implications for the accurate interpretation of array data, especially since a typical experimental design for Illumina Methylation arrays does not include technical or biological replication.

In this study, we present a series of comprehensive analyses that explore yet unidentified factors affecting the validity of Illumina Methylation array data. Paired longitudinal data from 142 paired blood samples (from 71 volunteers) collected 6 weeks apart was generated with the MethylationEPIC BeadChip v1.0, which enabled us to distinguish inter-individual DNAme variability with intra-individual DNAme data over time. Our results reveal new insights into factors affecting the variability of DNAme derived from the EPIC array and we thus propose a novel, data-driven method for the assessment of probe reliability. We expect that these findings will further improve existing pre-processing pipelines and the subsequent interpretation of next-generation Illumina Methylation array results.

## MATERIAL AND METHODS

### Sample collection and DNAme profiling in the clinical intervention study

93 individuals were recruited to the TACT (Turmeric-Anti-Inflammatory & Cell Damage Trial – clinical trial number NCT02815475) for a 6-week intervention study. There were 3 arms to the study: one group (‘Turmeric Capsule’ group; 25 patients: 17 females, 8 males) received a 400 mg Turmeric capsule providing 0.27g curcuminoids/day, a second group (‘Placebo’ Group; 24 patients: 16 females, 8 males) a sugar placebo (xylitol), and a third group were asked to regularly cook with Turmeric powder (‘Turmeric Powder’ group; 22 patients: 20 females, 2 males) providing 0.24g curcuminoids/day in their food every alternate day, all for a period of 6 weeks. Ethical approval number 16-WAT-23 was granted by Newcastle University’s SAgE ethics committee. 71 participants (53 females and 18 males) completed the study and provided full sets of usable 12-h fasting whole blood samples, which were collected at the start and end of the 6-week intervention into PAXGene DNA blood tubes (Becton Dickenson, 761165). Full blood counts were complemented with measurements of lymphocyte subsets (T/B/NK cells) using the fluorescently labelled antibodies CD3, CD4, CD8, CD19, and CD56, targeting T cells, T helper cells, cytotoxic T cells, B lymphocytes and NK cells (CD3 negative), respectively. Briefly, 50 μL blood were added to the antibody mix and incubated for 20’ at RT in the dark. To lyse the red blood cells, the mixture was incubated 10’ with 3 mL lysis buffer, washed (PBS-1%FBS) and cells were resuspended prior to flow cytometry analysis using a FACSCanto II (Becton Dickinson). DNA from the blood was extracted using the Machery Nagel NucleoMag® Blood 200 μL extraction kit (cat, 744501) and 500 ng total DNA was bisulfite modified using the EZ-96 DNA Methylation-Lightning kit (Zymo Research Corp, cat, D5047). 8 μl of modified DNA was subjected to methylation analysis on the Illumina Infinium MethylationEPIC BeadChip (Illumina, CA, USA) at UCL Genomics according to the manufacturer’s standard protocol.

### Normalization of MethylationEPIC data and immune cell subtype inference

Downstream analyses of the TACT study utilized raw β-values, obtained by formula (1) with raw intensities, as well as normalized β-values from three distinct pipelines: *minfi preprocessFunnorm* (10), *ChAMP* (11), and *ENmix* (12). β-values were regressed against the FACS-measured neutrophile and lymphocyte cell fractions for the first and second visits separately, and Infinium probes were considered cell type-dependent when FDR < 0.05 for both p-values associated with the slope of two linear regressions.

### Methylation changes linked to the clinical intervention

Two approaches were used to investigate differential methylation between two visits across the three treatment groups in the TACT study, either considering absolute differences in the original β-values between visits or considering differences in residuals from linear regression models (β-values versus real neutrophile cell fraction, the largest blood cell fraction) fitted on samples from the first visit only and then applied to all samples. Pairwise comparisons of the treatment groups were done using the Wilcoxon test, as well as a common comparison of all three groups using a Kruskal-Wallis test. Since the proportion of males in the ‘Turmeric Powder’ group was lower than in the other two groups, all tests were repeated on the female samples only.

### SNP analysis

Single nucleotide polymorphisms (SNPs) were identified from probes with underlying genetic sequence variation at target CpG sites listed by Pidsley et al. (Supplementary Table S4 in (5)). SNPs affect methylation profiles in specific ways depending on the position of the SNP relative to the target site. We defined the ‘SNP-II-0-effect’ associated with a 0-position (C base of target CG pair) of a Infinium Type II probe, which can cause false M (if SNP is G base) or false U signals (if SNP is T or A base), and the ‘SNP-II-1-effect’ associated with a 1-position (G base of target CG pair) of a type II probe, which may cause degradation of the total signal intensity (see Supplementary Figure S1). The ‘SNP-II-0-effect’ results in a tri-modal distribution of β-values of the type II probe where each mode is represented by carriers of one of three variants: CC – C on both chromosomes, C(SNP) or (SNP)C – C only on one chromosome, or (SNP)(SNP) for NON C on both chromosomes. Conversely, the ‘SNP-II-1-effect’ results in a tri-modal distribution of intensity levels of the type II probe, where each mode is represented by carriers of one of three variants: highest level for GG – G on both chromosomes, middle level for G(SNP) or (SNP)G – C only on one chromosome, and lowest level for (SNP)(SNP) for NON G on both chromosomes. On the level of β-values, the ‘SNP-II-1-effect’ results in a bi-modal distribution, with one mode corresponding to the (SNP)(SNP) variant and the second mode to the other two variants. Notably, SNPs in the other closest positions towards the end of the probe (likely 2-5 bp away) or large inserts and deletions in more distant positions can have the same ‘SNP-II-1-effect’. For type I-Red and I-Green probes, more SNP cases are possible that ultimately result in the same effects as described for type II probes, either a tri-modal β-value distribution (∼’SNP-II-0-effect’), a tri-modal intensity distribution (∼’SNP-II-1-effect’), or a combination of both (see Supplementary Figure S2).

### MI score calculation

We calculate the Mean normalized Intensity of each Infinium type probe (MI score) as follows:

For n samples, calculated across all type II or type I-Red or type I-Green probes separately:

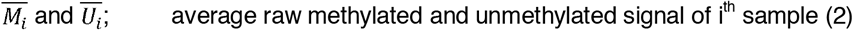

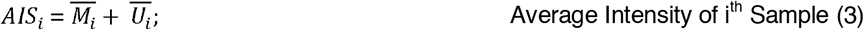

For each i^th^ sample and each j^th^ type II or type I-Red or type I-Green probe:

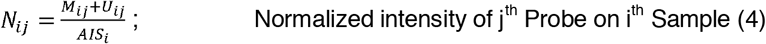

For each j^th^ type II or type I-Red or type I-Green probe:

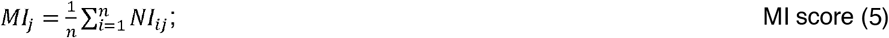

### Unreliability score calculation

First, intensities recorded in the Green and Red channels of the negative control probes on each array are collected (Green and Red noise, respectively) to create a Reliability Map (RM) for each probe type/colour separately, i.e., RM-II, RM-I-Green, RM-I-Red (see also Figure 2d). Each RM is a grid of pairs of fixed methylated and unmethylated values M_k_, U_I_ with k, l = (0, 5000, step = 100). For each pair of M_k_ and U_I_ in a RM, noise values are randomly selected 1000x and methylated noise M_err(k,I)_ and unmethylated noise U_err(k,I)_ defined as follows, for m = 1:1000:

- for type II probes:

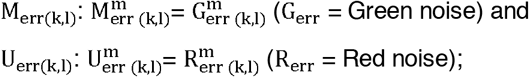

- for type I-Green probes:

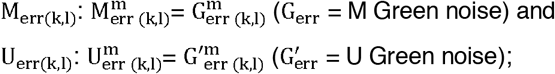

- for type I-Red probes:

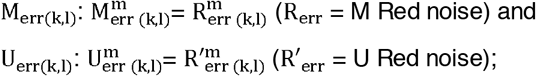

For each pair k and l we generate the artificial distribution of β-values by repeatedly adding 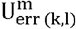 and 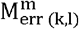 noise values to M_k_ and U_I_ respectively:

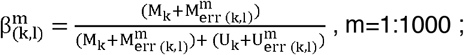

Then we calculate a Q score for a given distribution:

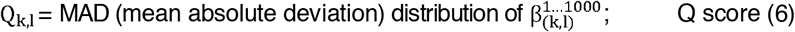

Thus, for each probe type the Reliability Map is a two-dimensional grid (G_k_, R_I_), of M and U signal intensities, where each cell is assigned the Q_k,I_ value, which is associated with the unreliability of the β-value obtained at the corresponding intensities. For each real (i.e., not modelled) pair of M_ij_ and U_ij_ signals from the i^th^ sample and each j^th^ probe average noise values are subtracted in the corresponding channels and the closest point (G*,R*) on RM and associated its Q*-value is retrieved to finally calculate the Unreliability score for each j^th^ probe across all n samples in the data set:

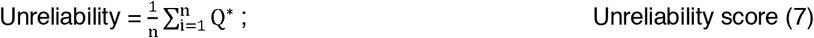

For intensity values outside the grid, then Q^*^ is assigned 0, that is very reliable.

### Unreliability and MI score dynamic thresholding

The relationship between unreliability and MI scores were examined for each probe type/colour separately, constructing smoothed curves using a generalized additive model (GAM). Because the dependence of Unreliability on MI rapidly decreases and then stabilizes after a so-called “critical point”, we propose a dynamic threshold for determining which probes are deemed unreliable in a given dataset, by determining the maximum of the second derivative of the smoothed.

### Unreliability and MI score validation

Two DNA methylation datasets comprised of true technical replicates were used to validate the utility of the unreliability and MI scores for probe reliability estimation. The first GSE174422 dataset with 128 duplicate pairs of female blood samples collected within a Sister Study and analyzed on an Illumina Infinium 450k Human Methylation Beadchip v1 (12) was downloaded from NCBI GEO. A second “Repeatability” dataset was generated in house using four technical replicates from the same DNA (2x) and bisulfite converted DNA mixtures (2x) obtained from three different sample types, i.e., fresh blood, frozen blood and cervical smears, from four female subjects that participated in the TirolGesund study (n = 3 x 4; see Supplementary Figure S3). Blood samples (2.5 mL) were stored in PAX gene blood DNA tubes (BD Biosciences) and DNA was isolated from fresh blood within a week after sample collection. The remaining blood was kept frozen at -20°C. DNA was additionally isolated from the frozen left-over samples and treated as a separate sample type. Cervical smears were collected and stored at room temperature with ThinPrep® Collection Kit (Hologic). Within a week after sample collection, cells sediments were transferred, washed (PBS) and pelleted (2,500 RPM, 10 min). Cell pellets were kept frozen at -80°C. DNA was isolated according to the tissue protocol of the Mag-Bind® Blood & Tissue DNA 96 Kit (#M6399-01, Omega Bio-tek) and quantified using the Quantifluor® dsDNA System (#E2670, Promega). 2 x 500 ng of DNA was bisulfite modified with the EZ DNA Methylation-Lightning kit (\#D5030, Zymo) and standardized to a concentration of 25 ng/uL BC-DNA. From each BC-DNA mixture 2 x 100 ng was processed on the Illumina Human MethylationEPIC v1.0 (#20042130) according to manufacturer’s instructions. To minimize batch effects, modified DNA from each sample type was processed randomly on array positions across two bead chips.

## RESULTS

### Unexpected DNAme variability in repeated blood samples

We initially studied whole blood DNA methylation profiles of 71 volunteers within the TACT study at two time points separated by a six-week interval (n=142). Although participants were allocated to one of three treatment groups (‘Placebo’, ‘Turmeric Capsule’ or ‘Turmeric Powder’), no significantly differentially methylated CpG sites (FDR > 0.05) were found with champ.dmp() for any of the four variants of β-values analyzed (raw or normalized with distinct published preprocessing pipelines; results not shown). For each CpG locus, we then calculated SD β, i.e., standard deviation of the β values within the population at visit 1, and Δβ, i.e., the average (over all individuals) of absolute differences in β values between visits (over time) for the same person. This revealed two distinct groups of CpGs targeted by Infinium probes in terms of DNAme variability (Fig. 1a): first, sites demonstrating a wide range of variability across the population of samples of a single visit (time point 1), and second, sites demonstrating a high degree of variability over time for the same individual. This distinction was evident both for both the raw and normalized β values.

**Fig. 1.**
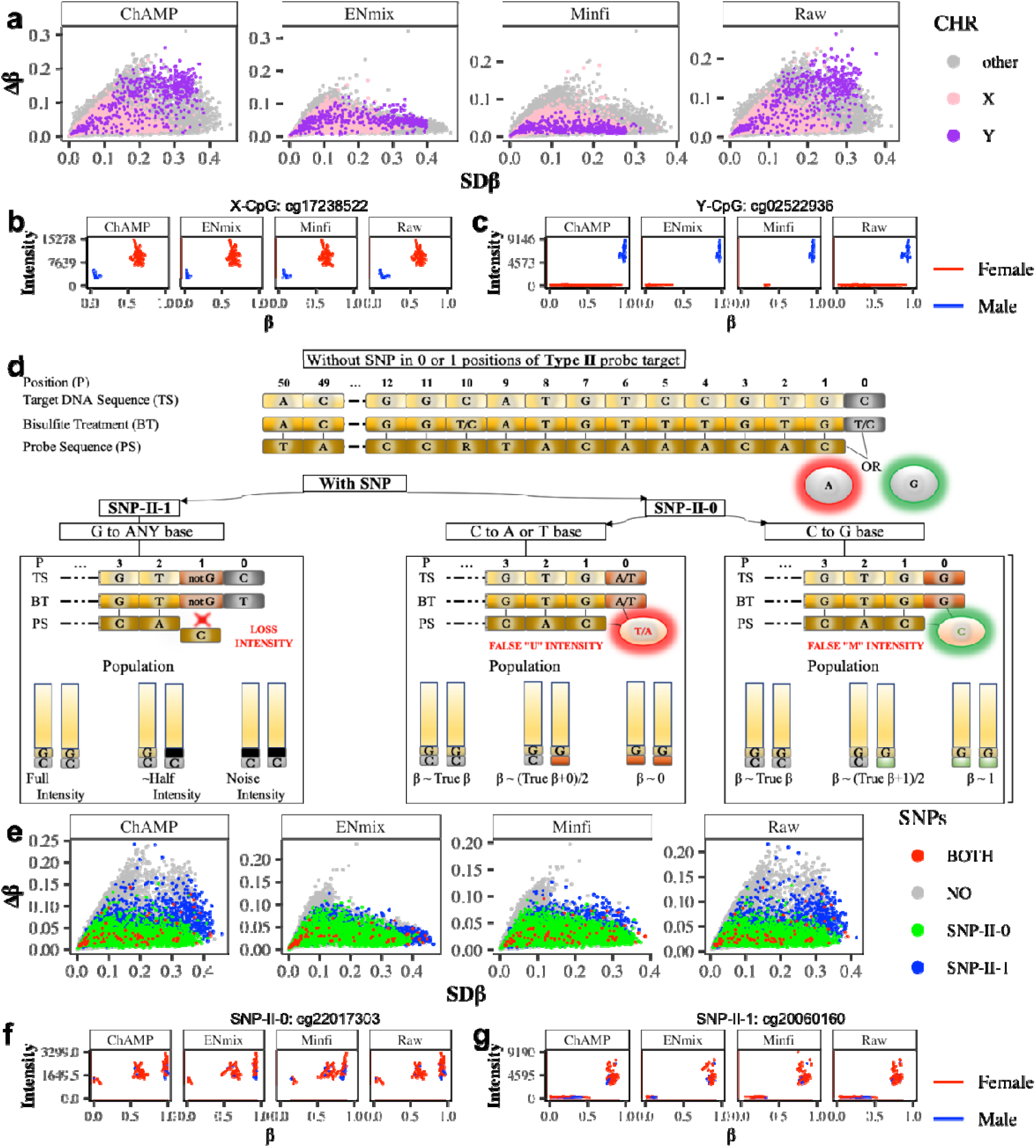
Variability associated with genetic factors (sex and genetic variants) in the TACT study. **(a)** Sex chromosome-associated probes demonstrate high variability both within population at visit 1 (SD β, x-axis) and over time (Δ β, y-axis). **(b)** Example of β value and intensity of an X-chromosome-associated probe in males and females. **(c)** Example of β value and intensity of a Y-chromosome-associated probe in males and females. **(d)** Two types of type II probes SNPs and their impact on signal intensity: SNPs in position 0 (SNP-II-0) result in false green or red signals, while SNPs in position 1 (SNP-II-1) result in a loss of signal (for type I probes see Supplementary Fig. 2a). **(e**) SNPs in position 0 and 1 both demonstrate high variability in the population at visit 1 (SD β), but SNPs in position 1 furthermore demonstrate high variability over time (Δ β; for type I probes Supplementary Fig. 2b). **(f)** Example of probe with 0-position SNP (SNP-II-0) shows tri-modal b-values distribution. **(g)** Example of probe with 1-position SNP (SNP-II-1) shows tri-modal intensities distributions.

**Fig. 2.**
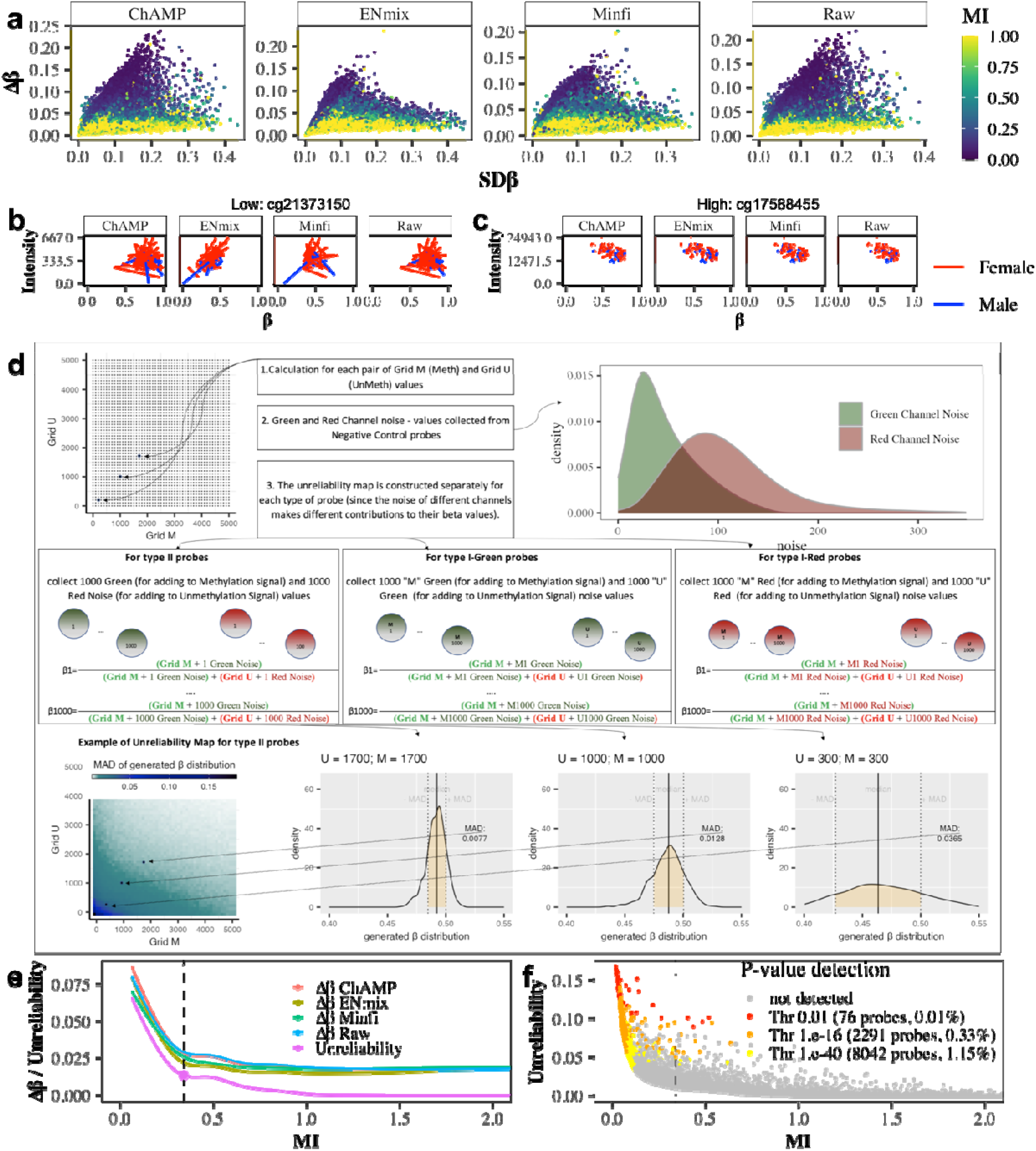
Impact of probe intensity on β value reliability in the TACT study. **(a)** Association of probe variability within participants at visit 1 (SD β) and over time (Δ β) with mean intensity (MI). Probes with low MI have high variability over time. Examples of a probe with low MI **(b)** or high MI **(c)**. Lines are connected by points corresponding to the individual in two different visits. β values from the same individual are closer for a probe with a high MI than on a probe with a low MI. **(d)** Reliability map of β values. **(e)** Smoothed curve shows the dependence of type II probe unreliability on mean intensity (MI). The point highlighted on the graph is the “critical point” marking a sharp change in dependence decline, which we recommend as a dynamic threshold for determining which probes are deemed unreliable (for type I probes and other data sets see Supplementary Figures S4, S5, S6). **(f)** Dependence of the type II probe unreliability on MI, highlighting probes which are detected using the p-value method at different threshold stringency (for type I probes and other datasets see Supplementary Figure S7).

Consequently, we checked to what degree the observed patterns of variability were linked to the genetic background of the targeted sites, in particular with sex chromosomes and single nucleotide polymorphisms (SNPs), sites typically removed or removed during preprocessing (13). When grouping probes by their chromosomal location (X, Y, and “other”), sex chromosome-associated probes within our dataset exhibited a high variability across the population, which is partially expected as our participant cohort included both men and women. However, strikingly, methylation values in probes targeting sex chromosomes also showed a high variability over time. This effect was less pronounced for *minfi* and *ENmix* normalized β values, likely due to the special normalization performed for sex chromosome-associated probes in these pipelines. As expected, M and U signal intensities at sex chromosomes was influenced by biological sex and sex chromosome copy number (Fig. 1b,c). The total signal intensity (M+U) of CpG probes mapping to the X chromosome is higher in females than males, since females have two X chromosomes. Conversely, the total signal intensity of CpG probes mapping to the Y chromosome is close to ‘0’, since females do not carry a Y chromosome, although some Y-CpGs might be in pseudoautosomal regions.

Infinium type I and II probes are based on inherently different designs, therefore we consider them separately for the remainder of our analyses. With respect to the Infinium type II probes, the two most relevant positional types of SNPs occur at position ‘0’ immediately after the 3’ end of the probe (SNP-II-0), where the SNP specifically affects the cytosine residue of the interrogated CpG, or at position ‘1’ at the very end of the 3’ end of the probe (SNP-II-1), where the SNP specifically affects the guanidine residue of the interrogated CpG (Fig. 1d). These two SNPs have distinct impacts on signal: a SNP-II-0 can result in false U or M signals, depending on the nature of the SNP replacing the ‘C’ residue, while a SNP-II-1 impairs hybridization and extension and results in a loss of signal. Other studies (5, 13) previously identified probes on the MethylationEPIC BeadChip v1.0 whose reliability is impacted by SNPs within sequence they target, and we have highlighted these probes in our dataset (see Fig. 1e for type II probes). Interestingly, the subset of probes with the highest Δβ, were neither a SNP-II-0 nor a SNP-II-1. Thus, like the sex-chromosome associated probes, SNP-associated probes contribute to a high degree of variability within the population, but they do not fully explain the high variability in DNAme data over time within individuals. We further found that SNP-II-1 resulted in bimodal β distributions (corresponding to signal or loss of signal; Fig. 1g), where the “true” variant is represented by the upper layer of intensity. SNP-II-0 that give rise to either false U or M signals yielded a trimodal β distributions (Fig. 1f), and the nature of the polymorphism, i.e., which base has replaced the cytosine residue, determined which particular mode corresponded to the values of the “true” (C/C) variant (see Supplementary Figure S1).

### Modelling the impact of signal intensity on β value reliability

Since the properties of both SNP-related and sex chromosome-associated probes are closely related to signal intensity (either through DNA quantity, false signals, or a loss of signal), but cannot fully explain the observed variability in repeated blood samples from the same individuals, we further scrutinized the impact of probe intensities on DNAme variability. For each probe on the array, we calculated a mean intensity (MI) value, which represents the corrected mean overall signal strength of the probe. Overall, probes with the highest level of time-dependent variability have a low MI value (Fig. 2a). Probes on the lower end of the MI scale in our dataset show a low reproducibility in paired blood samples (example cg21373150; Fig. 2b), whereas probes on the higher end of the MI scale a high reproducibility (cg17588455; Fig. 2c). We hypothesize that this high variability at low intensity levels is caused by a relatively higher impact of signal to noise. Therefore, the MI score may potentially allow for the identification of ‘noisy’ or ‘unreliable’ probes, i.e., probes which do not yield consistent β-values between two timepoints or replicates.

We created a simulation model to estimate the impact of noise on each probe’s “unreliability” by collecting the background intensities recorded by the negative control probes on the array and repeatedly adding methylated and unmethylated noise values, M-noise and U-noise, to a fixed grid of M and U signal pairs (Fig 2d, see Material and Methods section for further details). The resulting reliability maps summarize for each probe type/colour the mean absolute deviation of the resulting β distributions for each point in the grid (Q score) and are then subsequently used to assign an unreliability score for each probe in the final dataset, by averaging the matching Q scores for the measured β values across all samples. Dynamic thresholding for defining unreliable probes in each dataset is then achieved by examining the dependence of the unreliability scores on MI and finding the critical point of a smoothed curve where the dependence of unreliability (mirroring Δβ estimates) on MI stabilizes (Fig. 2e; Supplementary Figures S4, S5, S6).

Compared to the popular p-value detection method (detP, 15) to remove outlier probes as implemented in the *minfi* package (10), we improve by modelling the effect of noise on beta values obtained at different intensity levels, rather than comparing total intensities (across all genomic position in every sample) with distribution of total intensities on negative control probes (which only allows to estimate the ‘distance’ of probe intensities from the noise intensity, but does not allow to estimate the influence of noise on the final beta values). Furthermore, by allowing for data-driven thresholding we detect more unreliable probes than the statistical outliers alone, even compared to a very stringent threshold of p = 1.e-40 (6) for detP (Fig. 2f,e; Supplementary Figure S7, S8, S9, S10, S11).

### Linking mean intensity with probe sequence composition, target sequence copy number and unreliability scores

Investigating probe composition to identify factors associated with reliability, we found that probes with a low MI score tend to have a lower C content and target sequences with a lower G content (Fig. 3a; Supplementary Figure S12). Stronger physical binding between G-C base pairs than A-T base pairs could result in an increase in bound targets and fluorescent signal. Furthermore, probes targeting island, shore, shelf, and open sea regions inherently differ in their CG content (Fig. 3b). Correspondingly, open sea region probes have a lower MI and higher unreliability scores than probes located in island regions (Fig. 3c, 3d). As previously shown, the raw signal intensity of the probe depends on the number of copies on the DNA (in its bisulfite notation) complementary to the full 50bp sequence or a long-length nested subsequence of the probe (13). Here we additionally show that mean intensities depend both on copy number for 3? nested subsequences of probes longer than 30 nucleotides and C content (Fig. 3e).

**Fig. 3.**
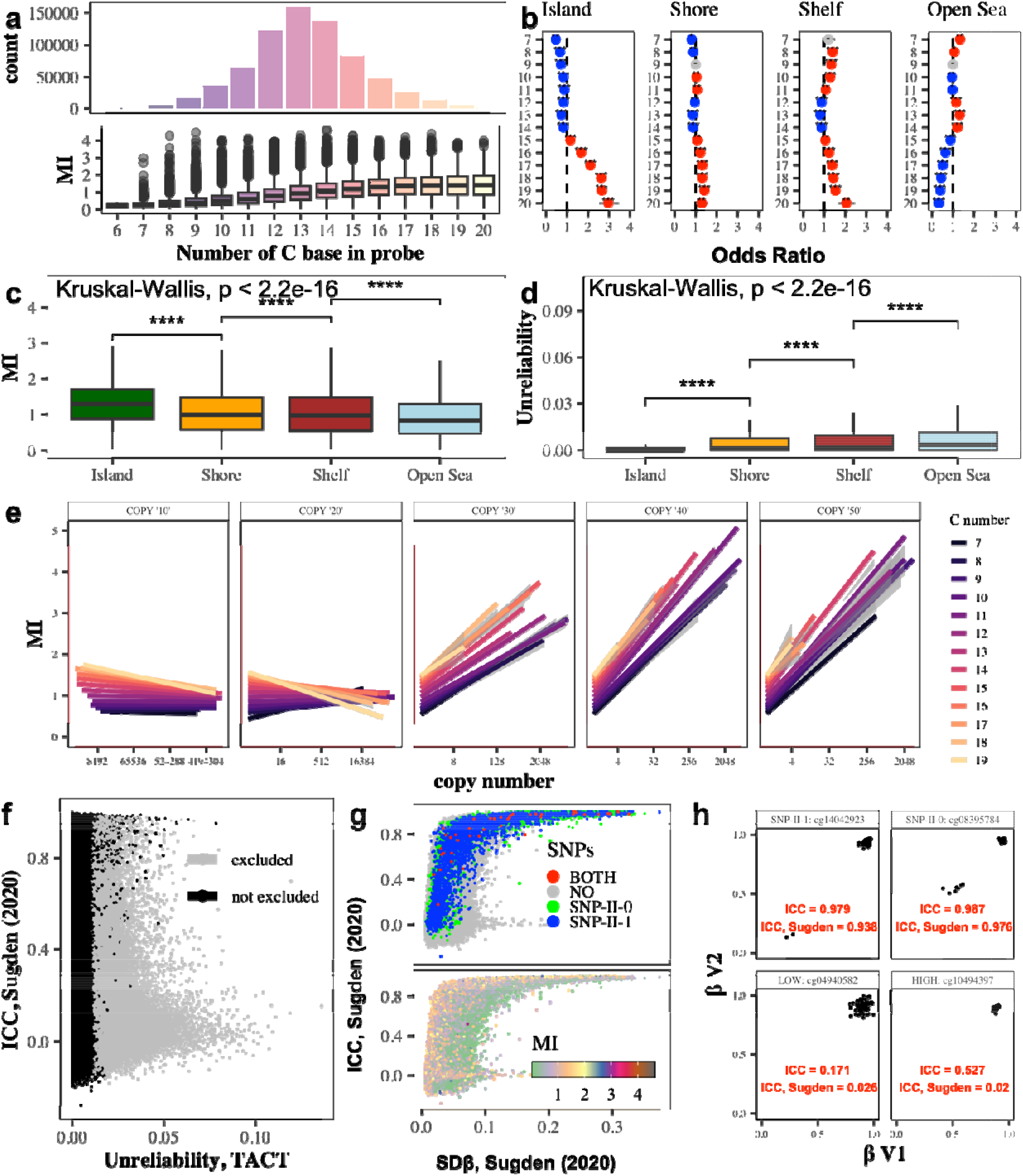
Type II probe features impacting signal intensity and reliability. **(a)** Dependence of mean signal intensity (MI) on C content of the probe (for type I probes and other data sets see Supplementary Figure S12). **(b)** Odds ratio for C enrichment in different DNA regions: Islands, due to their inherently high CG content, have a strong enrichment of probes with a large C content, which leads to the fact that on average MIs are higher **(c)** and Unreliability scores **(d)** are lower for Islands probes than for probes in other regions. **(e)** Dependence of the mean signal intensity (Y-axis) on raw signal intensity (X-axis) and C content of the probe sequence (color). **(f)** Reliability scores proposed by Sugden *et al*. (14) versus Unreliability scores defined here. **(g)** Association of Reliability scores proposed by Sugden *et al*. (14) with SNPs and MI. **(h)** Methylation status (β) and intraclass correlation coefficient (ICC) for example SNP-associated CpGs and probes yielding low signal intensity of probes.

Interestingly, our Unreliability score was not associated with a reliability measure proposed by Sugden *et al*. (14), which was calculated using ICC (Intraclass Correlation Coefficient) on β-values of repeat measurements of the same DNA samples (Fig. 3f). Furthermore, the reliability score from Sugden *et al*. does not correlate with MI (Fig. 3g – lower panel), and SNP-associated probes were deemed the most reliable using this measure (Fig. 3g – upper panel), which seems to be due to the high spread of β values in SNP 0 and SNP 1 (Fig. 3h – upper panels). Of note, some probes which have a similar Sugden reliability score had different ICC, MI and unreliability in our TACT data (Fig. 3h – lower panels). In contrast to the method proposed by Sugden *et al*., our method of assessing probe reliability is not based on cross-correlation of samples (which can be different in intensity, and therefore result in β bias), but instead offers insights into the reliability of probes based purely on intensity and noise distribution.

### Consistency of unreliability scores and probe MI with respect to biological and technical variation

Our method to estimate unreliability of Infinium probes is based on the analytical modelling of the effect of noise on probe intensities and explains the high values of Δβ well. However, since our dataset is comprised of paired biological replicate samples, we further investigate whether changes in methylation values over time could still be explained partially by biological influence, despite the absence of a treatment effect, and not only probe unreliability. Fractions of cell subtypes in the blood can change rapidly, for example the proportion of lymphocytes in peripheral blood can rapidly increase due to acute illness, influencing the β values that integrate the methylation status from all cell subtypes in the samples, and further explaining the variation seen in time for the same individuals. Indeed, the proportion of the two main immune blood cell subtypes, neutrophils, and lymphocytes, changed between the two visits (see example for type II probes in Supplementary Fig. S13a, S13b). We therefore selected probes whose β values are strongly influenced by these two immune subtypes (see Supplementary Fig. S13c) and evaluated the variability in these cell type-dependent probes between patients at visit 1 (SD β) and over time (Δ β; see Supplementary Fig. S13d).

To further demonstrate the consistency of the unreliability scores on true technical replicates, we analyzed two additional data sets: a published Illumina Infinium 450k Human Methylation Beadchip v1 dataset GSE174422 from 128 duplicate female blood samples, and a new EPIC v1.0 “Repeatability” dataset generated for this study from 4 quadruplicate fresh blood, 4 quadruplicate frozen blood and 4 quadruplicate cervical smear samples. Using raw β-values, we estimated variability by mean, absolute beta differences (Δβ) for each probe, confirming that variability increased with increasing unreliability scores and decreased with MI in the datasets with technical replicates (Figure 4a, 4b; Supplementary Fig. S14a, S14b, S15a, S15b). Also, for the technical replicate datasets, we detect more unreliable probes than the statistical outliers alone detected by the detP method with different thresholds and remove a significant portion of the lower variable probes deemed unreliable (Fig.4c; Supplementary Figures S8, S9, S10, S11). Furthermore, MI scores, which we use for dynamic thresholding in our unreliability method, correlate well across datasets (Fig. 4d), despite marked differences in total signal intensities for samples in the different datasets (Fig. 4e) and their noise distributions recorded from the negative control probes (Fig. 4f).

**Fig. 4.**
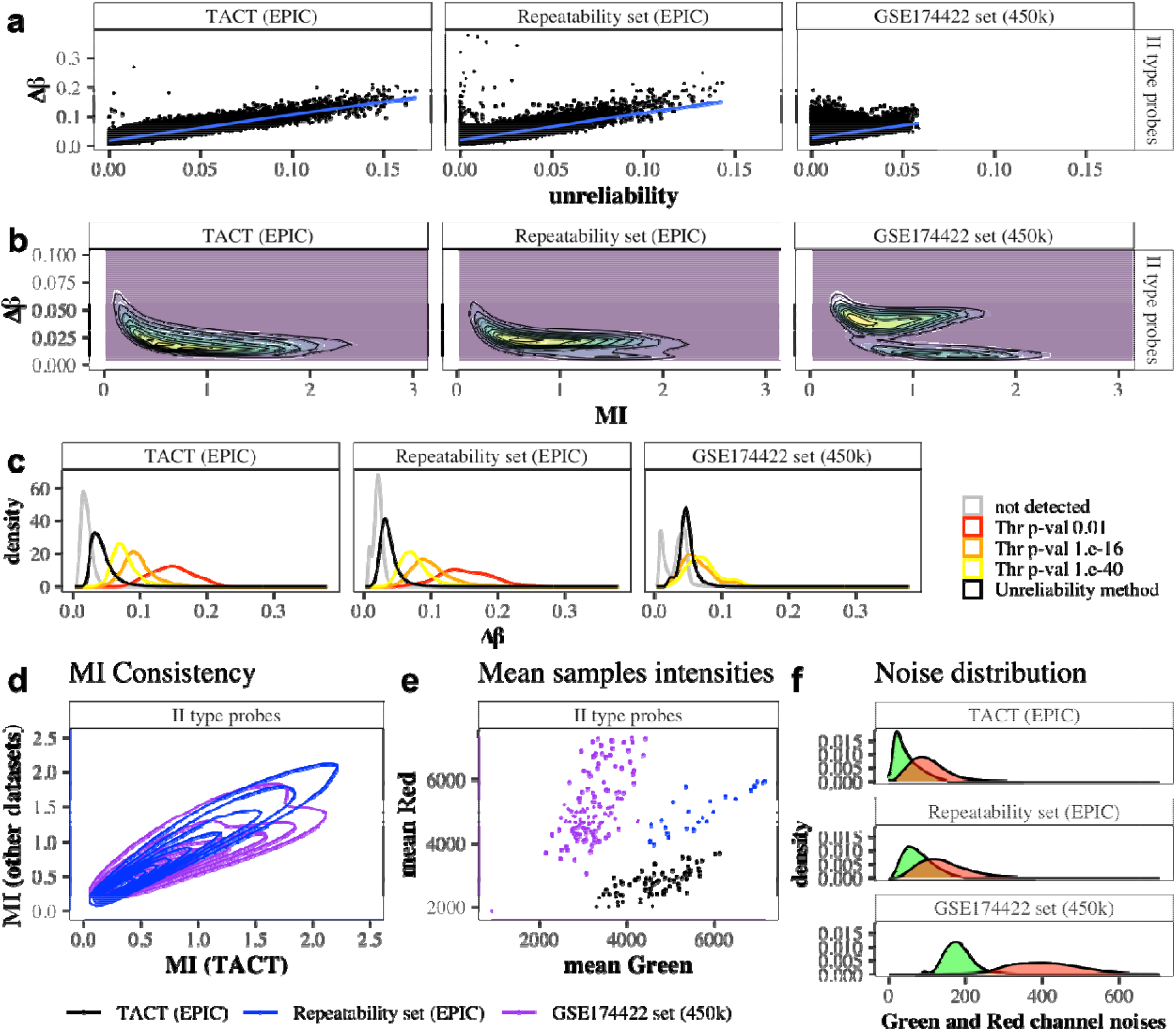
Validation of the unreliability method on datasets with technical replicates. Association of type II probe variability estimated by the averaged, absolute methylation differences between repeated samples (Δβ) with **(a)** the unreliability score and **(b)** the mean intensity (MI) in the TACT study (n=2x71 longitudinal paired blood samples), the Repeatability set (n=4x4x4 technical replicates for fresh blood, frozen blood and cervical smear samples) and GSE174422 (n=2x128 technical replicates for blood samples). For type I probes see Supplementary Fig. S14a, S14b, S15a, S15b. **(c)** Distribution of Δβ for all type II probes (grey) or those removed by the p-value detection method at different threshold settings (red, orange, yellow) and the Unreliability method (black) in the different data sets. (**d)** Correlation of MI measured for type II probes across data sets. **(e)** Mean red versus green total signal intensities of the samples in the different data sets. **(f)** Noise distributions obtained from the negative control probes in the different datasets.

### Implications for the newer MethylationEPIC BeadChip v2.0

A new MethylationEPIC BeadChip v2.0 was released in November 2022. The new manifest reports that ‘underperforming’ probes were removed compared to the v1.0 manifest (approximately 140,000, i.e., 23% of all type I probes and 15% of all type II probes). However, we found no evidence of an enrichment for probes with high unreliability or low MI in those discarded for v2.0 (Supplementary Figure S16). Therefore, we assume that the issues raised here will remain of high importance also for the newer version of the EPIC array. We also note that despite the announced large-scale removal of SNPs, some SNP 0 (∼15%) and 1 (∼25%) probes remained on v2.0. In addition, we observed that some probes are not marked as containing SNPs (neither by Pidsley (5) and Zhou lists (13), nor by the Illumina Manifest), but clearly demonstrate SNP-II-0 or SNP-II-1 behaviour (Supplementary Figure S17, S18).

## DISCUSSION

Despite considerable investment in improving existing analysis pipelines for popular Illumina methylation arrays, room for improvement remains. To facilitate the generation of meaningful findings that will not only increase our understanding of the epigenome and its relationship with health and disease, but also translate into clinically useful tools, it is vital that we fully understand how robust these DNA methylation arrays perform. Here we show that DNAme data from paired whole blood samples taken from the same individuals display variability over time which cannot be attributed to underlying genetic or biological factors alone. Much of the ‘unexplained’ temporal variability in the current study can be attributed to probe quality, which is primarily dictated by the probe sequence complexity and genome location.

Noise affects methylation values differently at different intensity levels: it has a dramatic effect on β-values at low intensity, while at high intensities, the signal cancels out the effect of noise. We therefore developed an approach for assessing the unreliability of β-values in a data-driven manner, using the negative control probes on the arrays to model the contribution of noise to any of the final signal intensities in a specific data set. Our new unreliability score correlates well increasing degrees of variability observed between repeated samples, both in longitudinal data set as well as in two validation sets with technical replicates. By modeling the noise distribution for each dataset for type II and type I/color probes independently, we were able to detect more unreliable probes compared to an existing detP method for detecting outlier probes ((15) with 0.01 p-value threshold, (16) with 1.e-16 p-value threshold, (6) with 1.e-40 p-value threshold) .

There is a marked difference in signal intensity and quality based on the C-content of the probe sequence and CpG content. On the one hand, this observation can help Illumina to achieve leveling of such an effect within the technological process, on the other hand, it will allow scientists to provide more qualitative comparisons on different regions of DNA, for example separating Islands and the Open Sea CpGs. Excluding low-intensity probes or unreliable from the analysis could help increase the detection of differentially methylated CpGs for different phenotypes and improve both the accuracy and precision of existing and emerging predictive models on this type of DNAme data. Beyond the mere exclusion of unreliable probes, new correction or normalization methods may emerge in the future based on the results of this work that could instead salvage the data generated from these probes.

Factors contributing to laboratory or methodological bias, such as sample storage and hybridization procedures, are relatively underexplored. Samples from different studies tend to be of different quality, yielding different average intensities depending on the instruments used and the exact laboratory protocols, which in turn can also affect the reproducibility of β-values on probes with different intensities and estimated probe reliability. Therefore, accounting for probe reliability and raw signal intensities during initial quality control may also improve the reproducibility of DNAme studies across laboratories.

In summary, we developed a new computational method to further refine existing preprocessing methods for Illumina methylation array data by excluding unreliable probes from downstream analyses. We implemented our methods to calculate MI and unreliability scores into an R package, epicMI, which is publicly available on GitHub (17).

## DATA AVAILABILITY

The raw methylation data from Repeatability set is available under controlled-access under the accession number EGAS00001007184 from the European Genome-phenome Archive (EGA), which is hosted by the EBI and the CRG. Raw DNA methylation data from the TACT study is not deposited to a public repository, because volunteers did not specifically consent to make data containing genetic information (that could identify individuals) publicly available. Preprocessed data with probes at SNP positions removed can be made available upon request from researchers in countries compliant with GDPR law only. The R code for calculation of the MI and unreliability scores is available in a GitHub repository (https://github.com/ChVav/epicMI) (17).

## ACKNOWLEDGEMENT

We thank all volunteers who provided samples for this research.

