## Supplementary Files for "Technical and biological sources of unreliability of Infinium probes on Illumina Methylation microarrays"

**a**

| | | | | TRUE FRACTIONS (when **C** in 0-position on 1 or 2 chromosomes) | | FALSE FRACTIONS (when **SNP** in 0-position on 1 or 2 chromosomes) | | |
|---|---|---|---|---|---|---|---|---|
| **TYPE** | **ALTERNATIVE** | **POSSIBLE VARIANTS** | **PROBABILITY OF VARIANTS** | **GREEN FRACTION** | **RED FRACTION** | **FALSE GREEN FRACTION** | **FALSE RED FRACTION** | **GREEN/(GREEN + RED)** |
| **NON $SNP_0$** | **[C]** | CC | P=1 | X | 1-X | 0 | 0 | BETA = X |
| **$SNP_0$** | **[C/G]** | CC | $P = P_C P_C$ | X | 1-X | 0 | 0 | BETA = X |
| | | CG or GC | $P = 2P_C P_G$ | 0.5X | 0.5(1-X) | 0.5 | 0 | BETA =0.5X + 0.5 |
| | | GG | $P = P_G P_G$ | 0 | 0 | 1 | 0 | BETA = 1 |
| | **[C/T]** | CC | $P = P_C P_C$ | X | 1-X | 0 | 0 | BETA = X |
| | | CT or TC | $P = 2P_C P_T$ | 0.5X | 0.5(1-X) | 0 | 0.5 | BETA =0.5X |
| | | TT | $P = P_T P_T$ | 0 | 0 | 0 | 1 | BETA = 0 |
| | **[C/A]** | CC | $P = P_C P_C$ | X | 1-X | 0 | 0 | BETA = X |
| | | CA or AC | $P = 2P_C P_A$ | 0.5X | 0.5(1-X) | 0 | 0.5 | BETA =0.5X |
| | | AA | $P = P_A P_A$ | 0 | 0 | 0 | 1 | BETA = 0 |

**b**

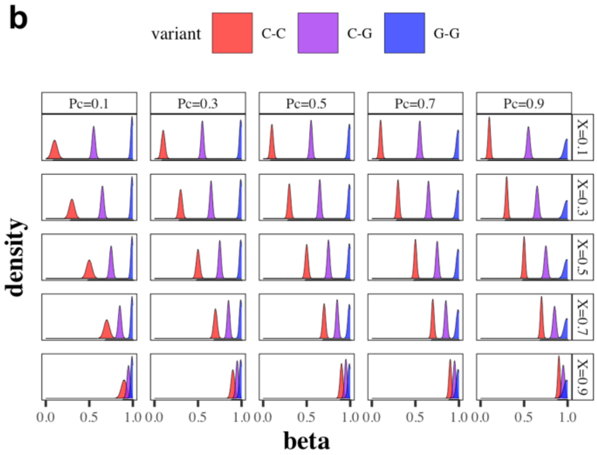


**c**

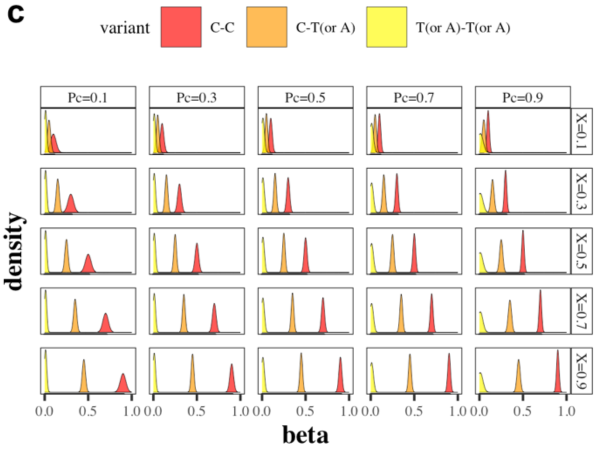


**Figure S1. Formation of tri-modal distributions of beta-values in the presence of a SNP in 0-position (C in the CG pair). (a)** The table shows how, in the case of the presence of SNP in the 0-position (in C base of target CG pair), depending on the nucleotide (G or T/A), a trimodal distribution of beta-values arises, where each mode is represented by carriers of a certain variant. For example, in the case when C base in the CG pair can be replaced by G in population, then the entire population is divided into three groups of carriers of different variants (CC - C base on both chromosomes; CG or GC - when C (or G) base is present only on one chromosome; GG - G base on both chromosomes). Carriers of the CC variant will receive true green and red signals (let their average beta value be X). Carriers of the CG or GC variant will receive true green and red methylation signals from one chromosome and a false green signal from the other (which will cause their beta-values to shift to the right, towards increased methylation and the average signal will be approximately 0.5X + 0.5); Carriers of the GG variant will get false green signal from both chromosomes and the average beta-values will be close to 1). Similarly, trimodal distributions of beta-values are formed if the C base in CG pair can be replaced in the population by T or A. In this case, for carriers of T or A bases (on one or both chromosomes) a false red signal will accumulate, and the modes of their beta-values will shift to the left (will be lower) than the mode of carriers of the CC variant. The mode density in such tri-modal distributions is determined by the probability of the variant, defined through the probability of the SNP or the probability of C. Simulation of tri-modal distributions for both situations: when C base in the target CG pair can be replaced by G base **(b)** or T/A base **(c)**, depending on the probability C and on X - true beta signal (from carriers of CC variant).

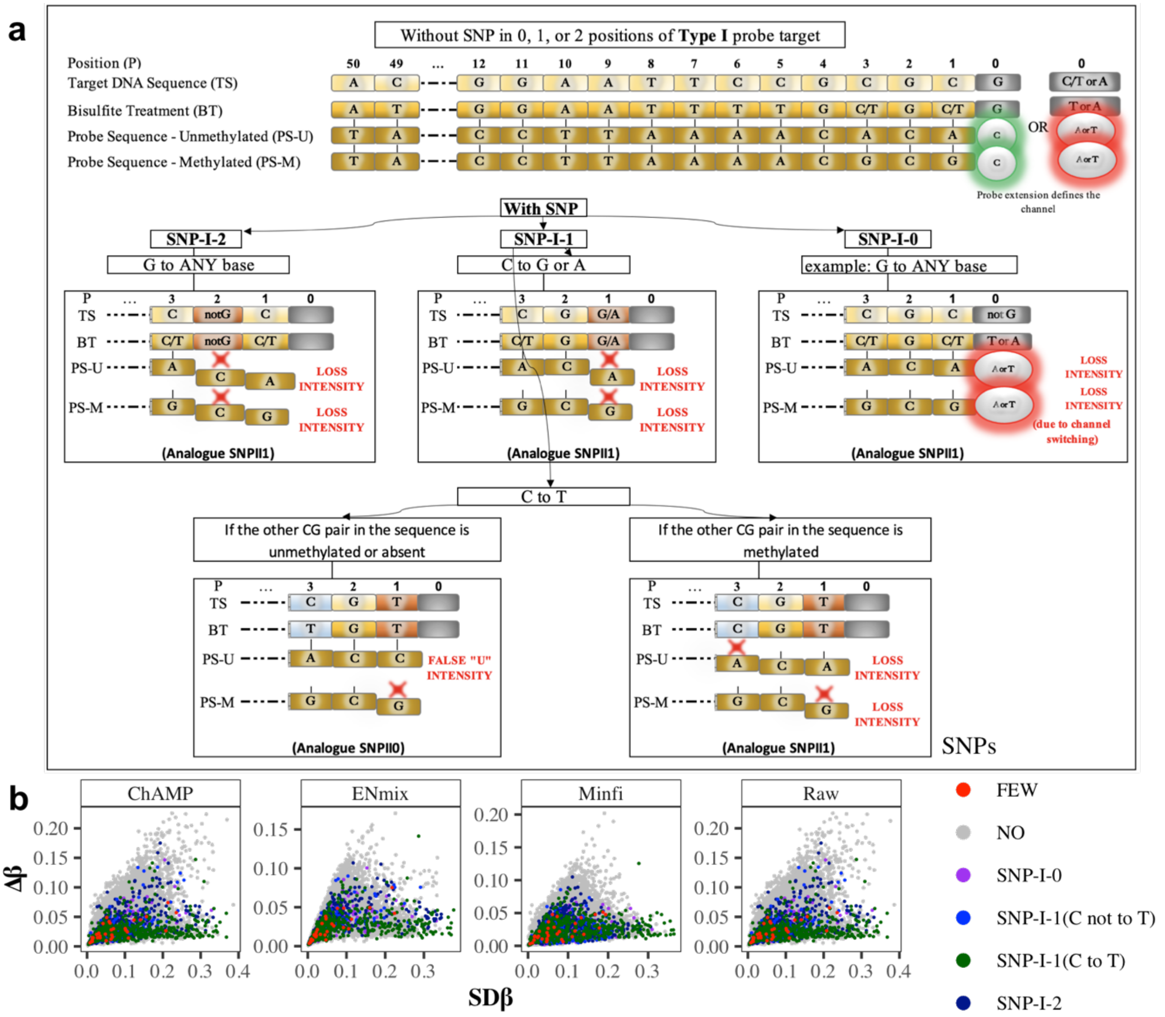


**Figure S2. Variability associated with genetic variants (Type I probes). (a)** Three types of SNPs and their impact on signal: SNPs in position 0 (SNP-I-0) can be result a loss of signal due to channel switching in case if original G base has as a variant any other base (as shown on example), or if original C/T/A base has as a variant G base; SNPs in position 2 (SNP-I-2) and 1 when C base base has as a variant G or A (SNP-I-1(C not to T)), or when C has a variant T (SNP-I-1(C to T)), and other CG pairs in the sequences are methylated result in a loss of signal due to the fact that the bisulfite sequence is no longer complementary to the U and M probes; SNPs in position 1 when C base has a variant T (SNP-I-1(C to T)) and other CG pairs in the sequences are unmethylated or absent result in false U signals. **(b)** SNPs in position 2,1 and 0 demonstrate high variability in the population at visit 1 (SD β) and SNP-I-0, SNP-I-2, SNP-I-1(C not to T) and part of SNP-I-1(C to T) show the same effect as SNP-II-1 probes (i.e. demonstrate high variability over time (Δ β)), and other part of SNP-I-1(C to T) shows the same effect as SNP-II-0 probes.

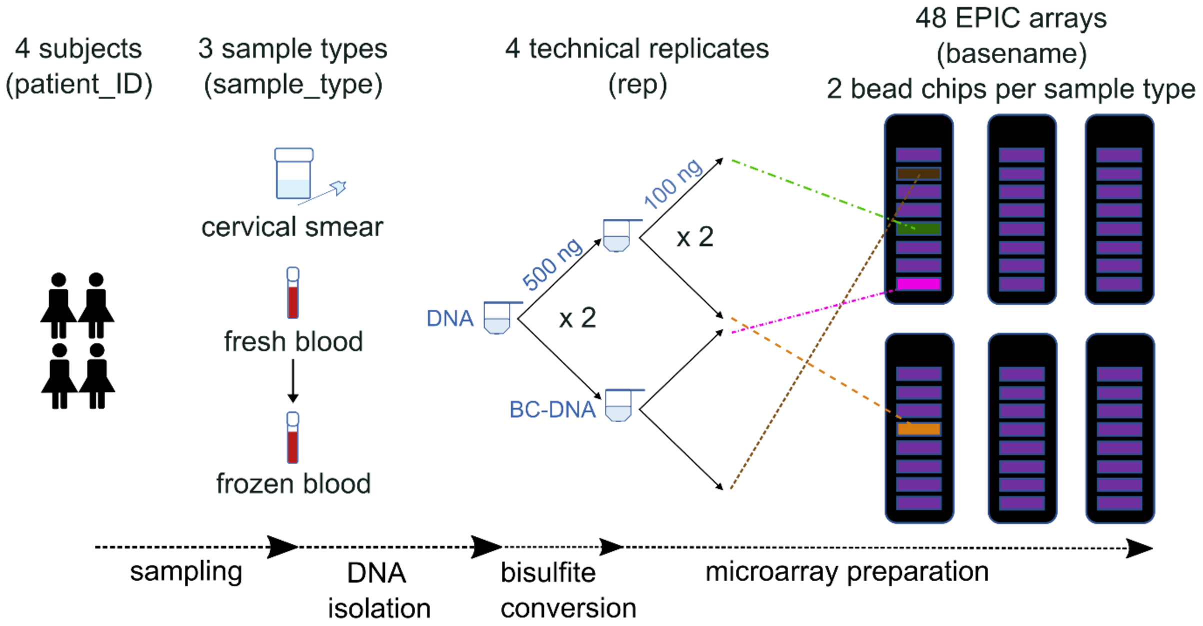


**Figure S3. Experimental design repeatability set.** Whole blood and cervical smear samples were collected from four female, healthy subjects. DNA from the blood was isolated either directly (fresh blood) or from the left-over sample that was kept frozen for prolonged storage. Four technical replicates were created from each DNA mixture: two times 500 ng from the same mixture was bisulfite converted (BC), and from each BC-DNA mixture two times 100 ng was prepared for hybridization to the microarrays. For each sample type, i.e. cervical smear, fresh blood or frozen blood, the positions of the samples and replicates were randomized across two bead chips.

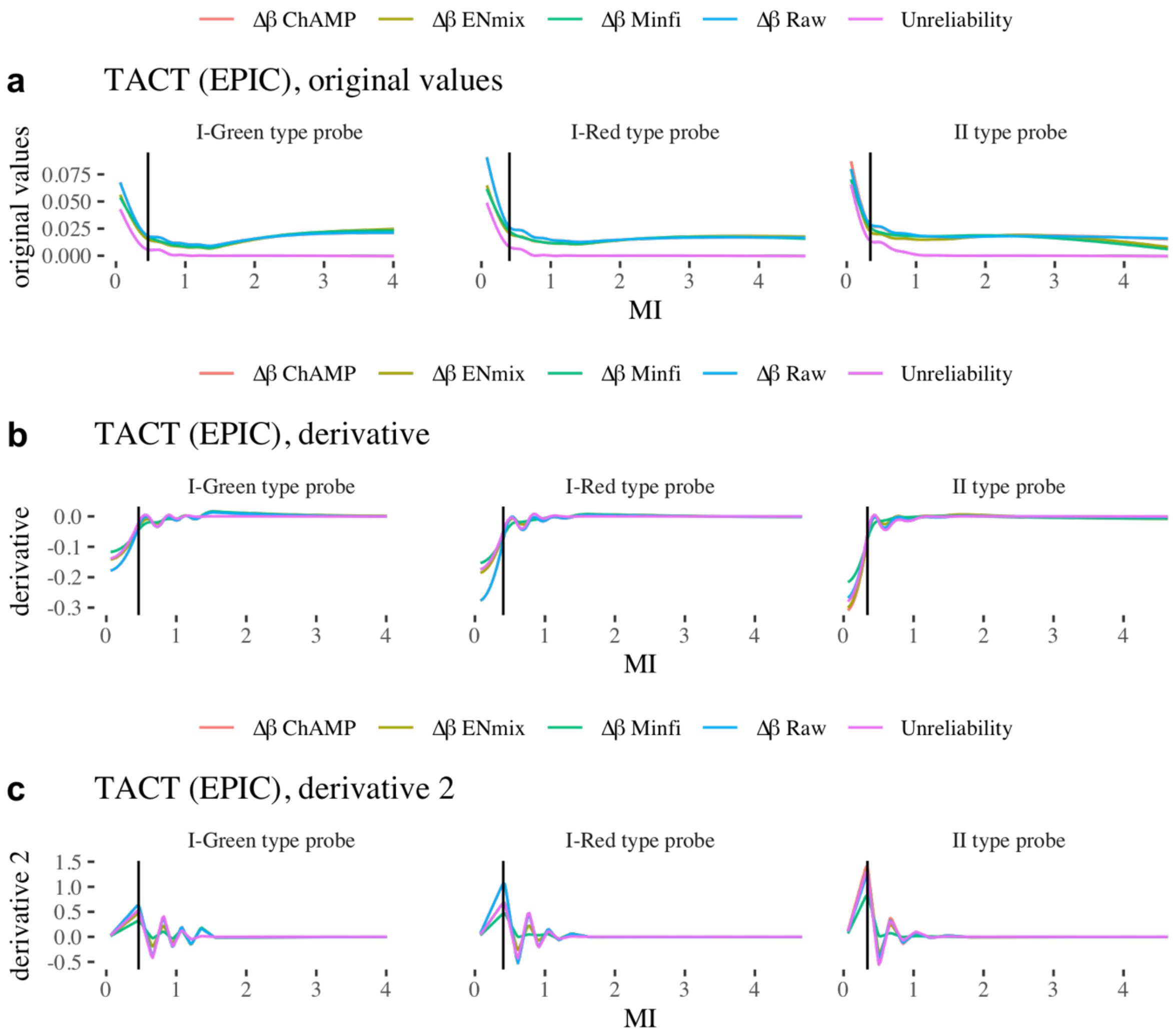


**Figure S4. (a)** Averaged, absolute methylation differences in methylation values between repeated samples (Δβ) and associated unreliability scores as a function of MI in the longitudinal TACT dataset with ('ChAMP', 'Enmix', 'Minfi') and without ('Raw') different normalization methods are plotted for each probe type/color, and **(b)** first and **(c)** second derivatives thereof.

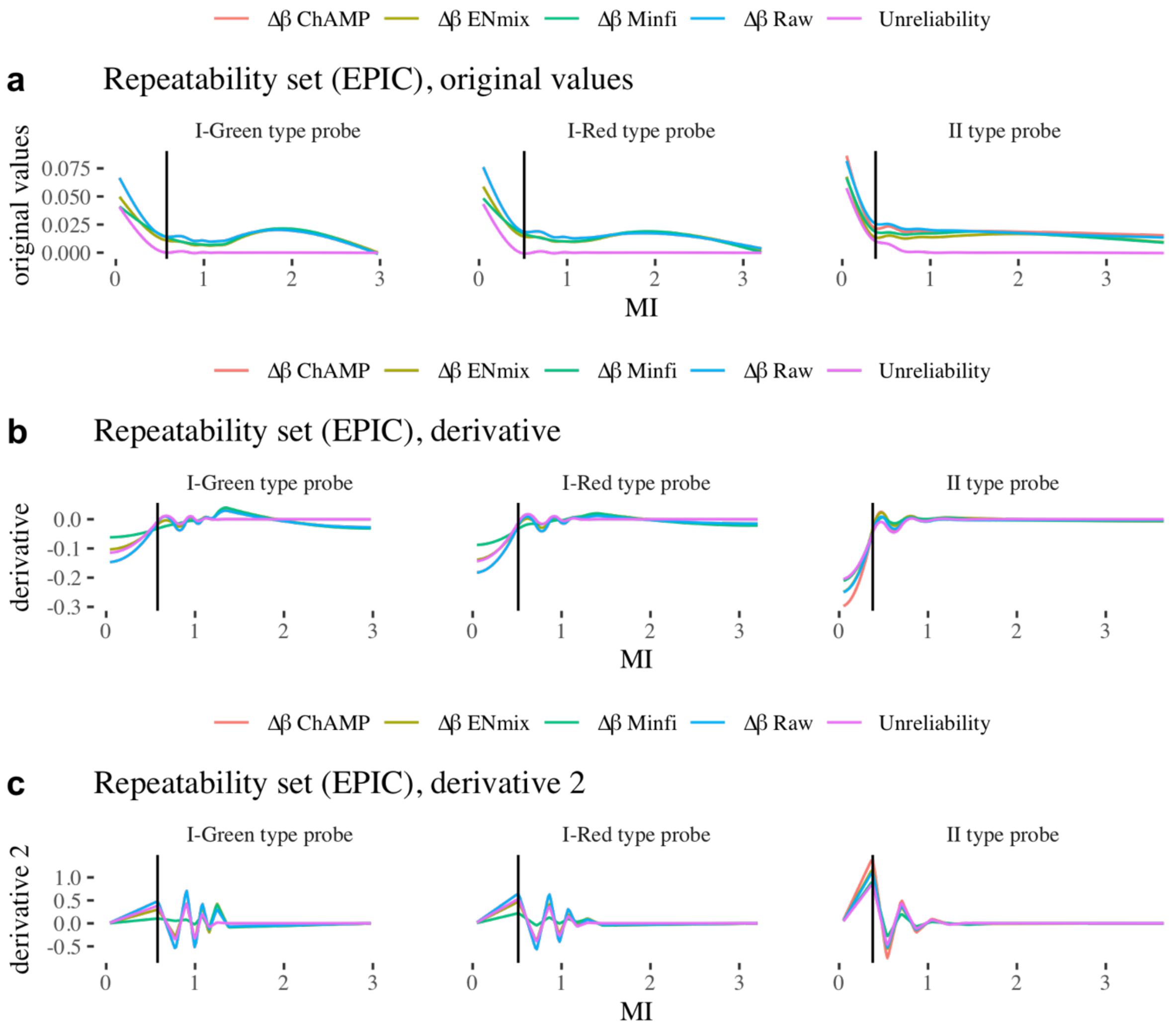


**Figure S5. (a)** Averaged, absolute methylation differences in methylation values between technical replicates (Δβ) and associated unreliability scores as a function of MI in the longitudinal Repeatability dataset with (‘ChAMP’, ‘Enmix’, ‘Minfi’) and without (‘Raw’) different normalization methods are plotted for each probe type/color, and **(b)** first and **(c)** second derivatives thereof.

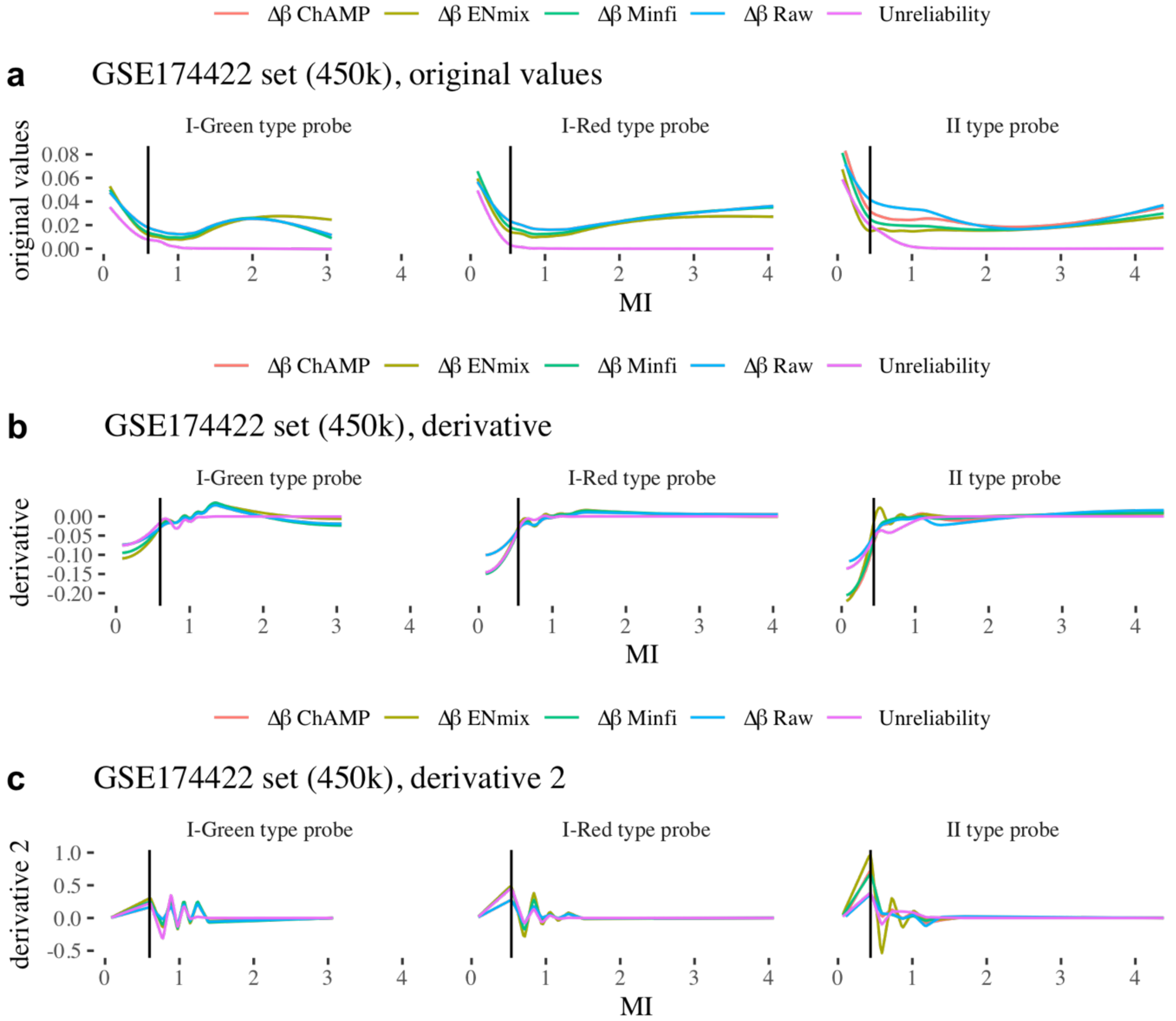


**Figure S6. (a)** Averaged, absolute methylation differences in methylation values between technical replicates (Δβ) and associated unreliability scores as a function of MI in the GSE174422 dataset with ('ChAMP', 'Enmix', 'Minfi') and without ('Raw') different normalization methods are plotted for each probe type/color, and **(b)** first and **(c)** second derivatives thereof.

**a**

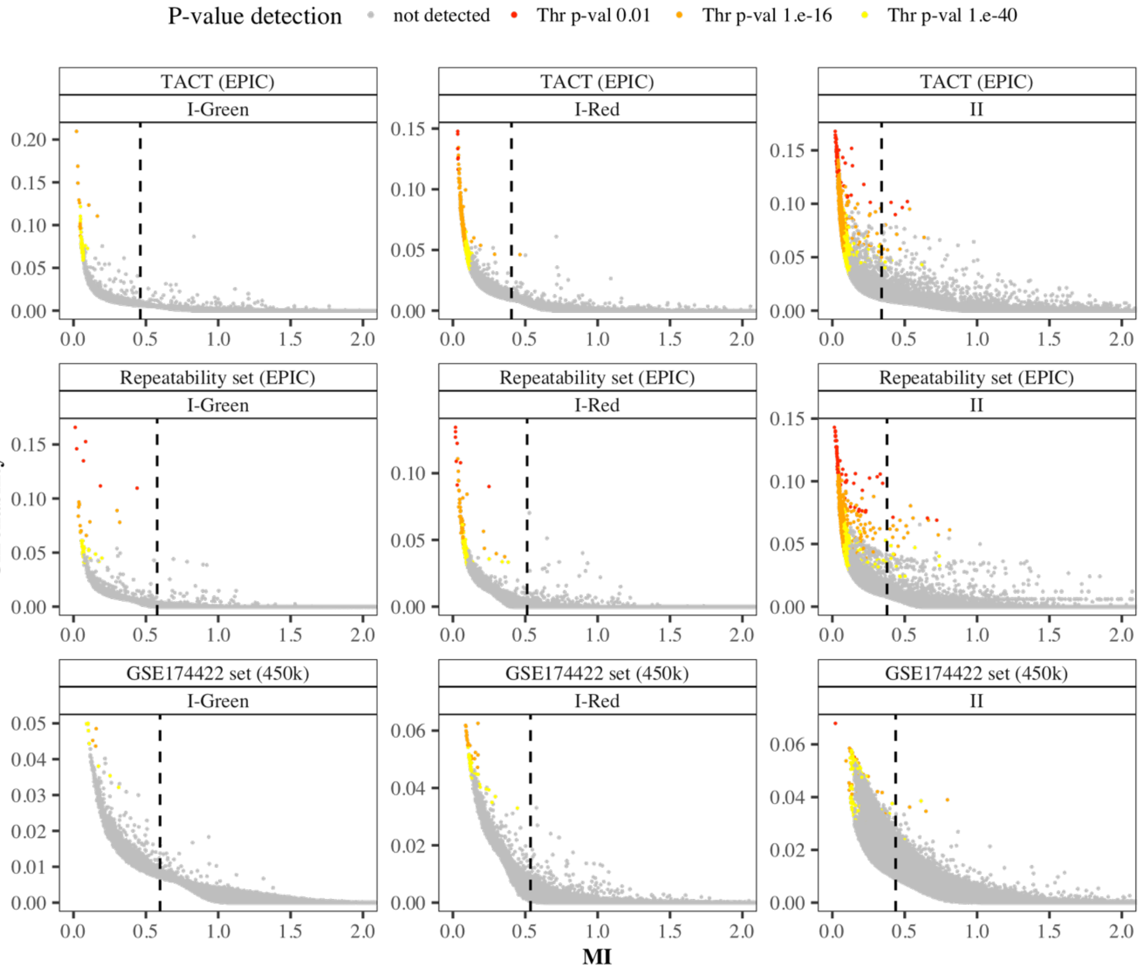


**b**

| Dataset | I-Green | | | | | | I-Red | | | | | | II | | | | | |
|---|---|---|---|---|---|---|---|---|---|---|---|---|---|---|---|---|---|---|
| | #probes | #numbes (without SNPs and X/Y CHR) | #probes detected with p-val 0.01 | #probes detected with p-val 1.e-16 | #probes detected with p-val 1.e-40 | #probes detected with Unreliability | #probes | #numbes (without SNPs and X/Y CHR) | #probes detected with p-val 0.01 | #probes detected with p-val 1.e-16 | #probes detected with p-val 1.e-40 | #probes detected with Unreliability | #probes | #numbes (without SNPs and X/Y CHR) | #probes detected with p-val 0.01 | #probes detected with p-val 1.e-16 | #probes detected with p-val 1.e-40 | #probes detected with Unreliability |
| TACT (EPIC) | 49939 | 48072 (100%) | 0 (0%) | 13 (0.03%) | 56 (0.12%) | **5684 (11.82%)** | 92198 | 88687 (100%) | 6 (0.01%) | 229 (0.26%) | 764 (0.86%) | **9967 (11.24%)** | 723722 | 697357 (100%) | 76 (0.01%) | 2291 (0.33%) | 8042 (1.15%) | **87889 (12.6%)** |
| Repeatability set (EPIC) | 49939 | 48072 (100%) | 6 (0.01%) | 18 (0.04%) | 47 (0.1%) | **5481 (11.4%)** | 92198 | 88687 (100%) | 8 (0.01%) | 59 (0.07%) | 181 (0.2%) | **10186 (11.49%)** | 723659 | 697297 (100%) | 61 (0.01%) | 899 (0.13%) | 3486 (0.5%) | **85016 (12.19%)** |
| GSE174422 set (450k) | 46289 | 44588 (100%) | 0 (0%) | 3 (0.01%) | 11 (0.02%) | **4925 (11.05%)** | 89187 | 85930 (100%) | 0 (0%) | 18 (0.02%) | 62 (0.07%) | **10199 (11.87%)** | 350036 | 335980 (100%) | 2 (0%) | 33 (0.01%) | 162 (0.05%) | **41796 (12.44%)** |

**Figure S7. (a)** Dependence of probe unreliability on MI, highlighting probes which are detected using the p-value method at different threshold stringency. **(b)** Number of probes removed by the respective methods.

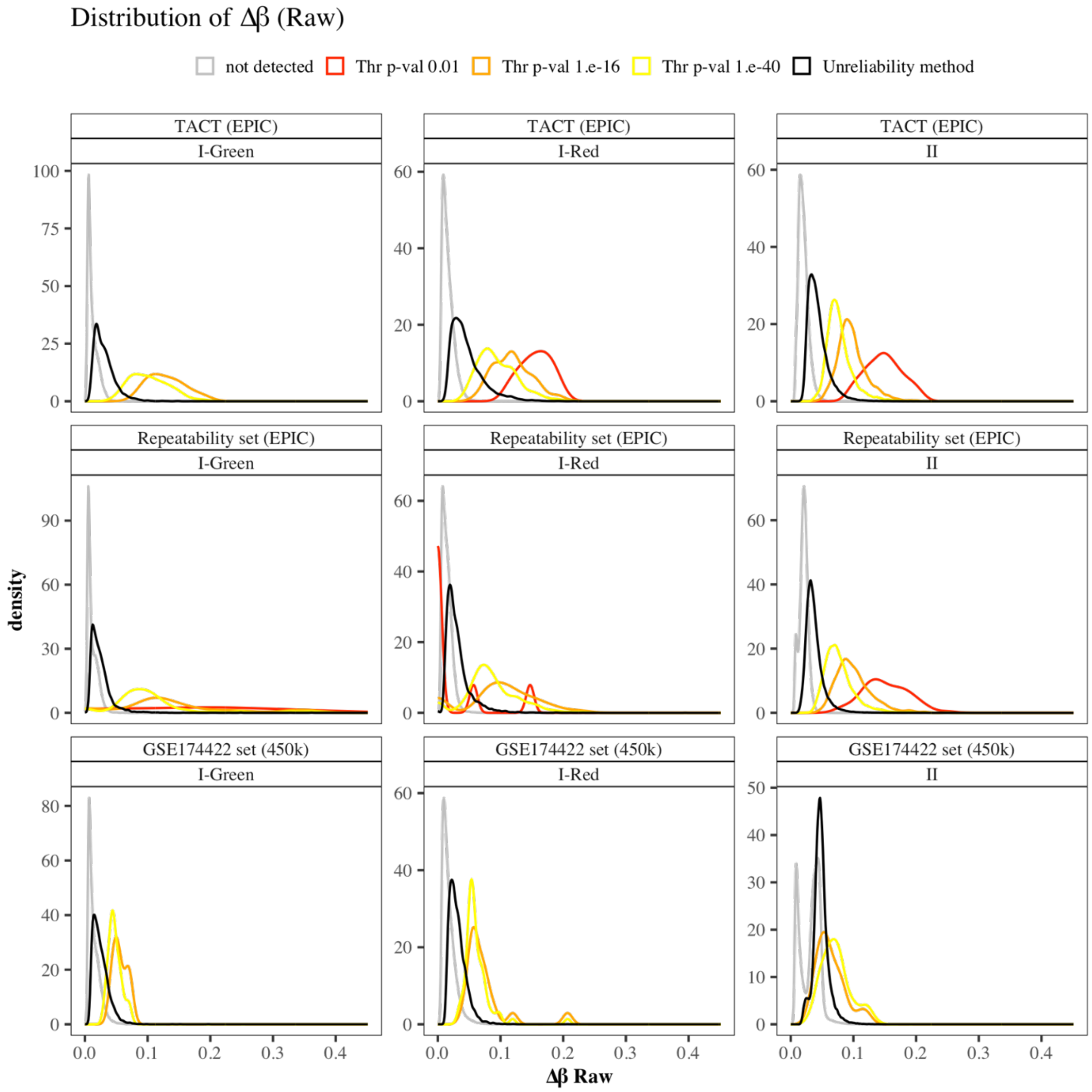


**Figure S8.** Distribution of the averaged, absolute methylation differences in methylation values between repeated samples and technical replicates (Δβ) for all probes (grey) or those removed by the p-value detection method at different threshold settings (red, orange, yellow) and the Unreliability method (black). β-values were not normalized.

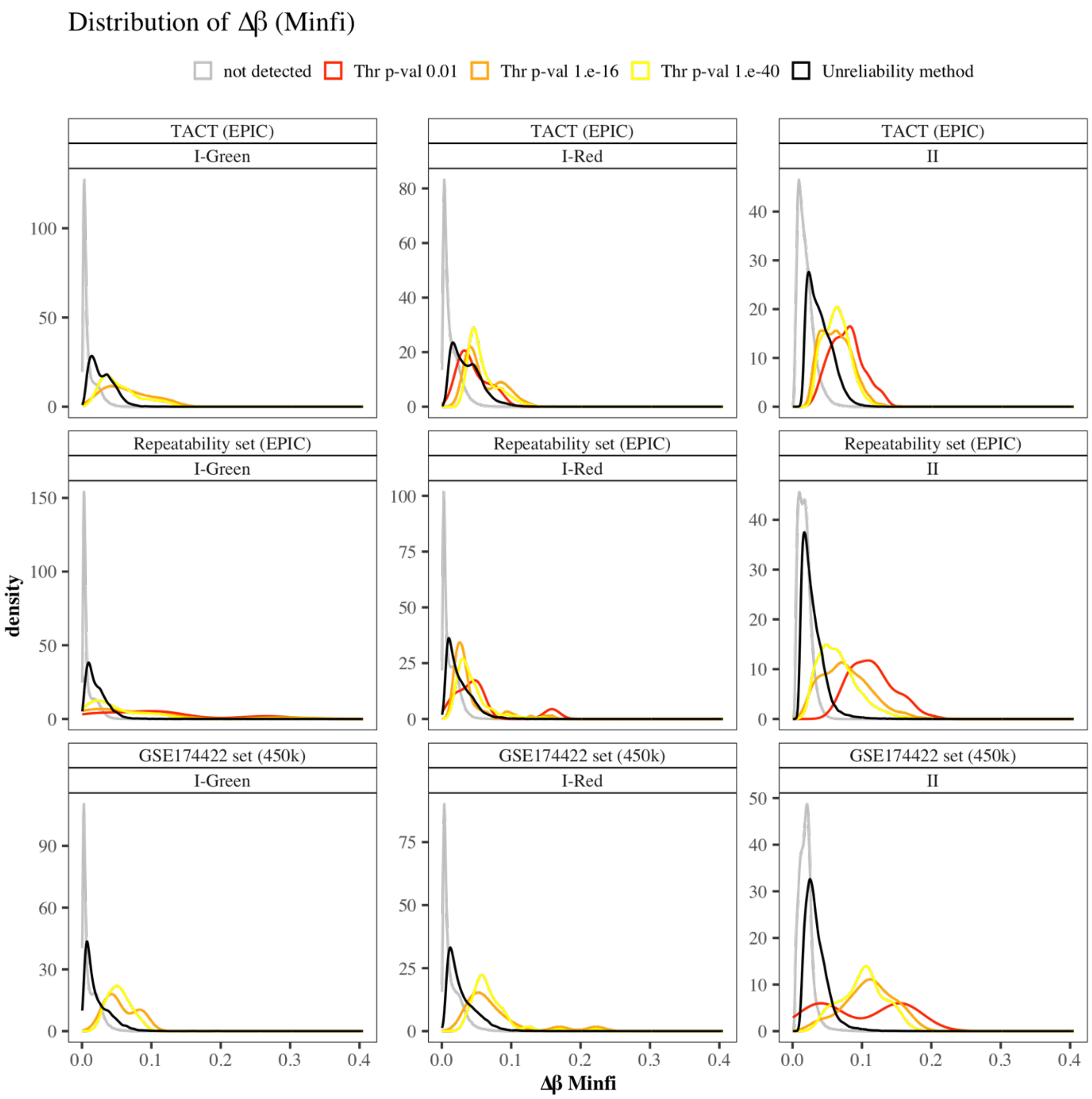


**Figure S9.** Distribution of the averaged, absolute methylation differences in methylation values between repeated samples and technical replicates (Δβ) for all probes (grey) or those removed by the p-value detection method at different threshold settings (red, orange, yellow) and the Unreliability method (black). β-values were normalized using the preprocessFunnorm() function in the R package *minfi*.

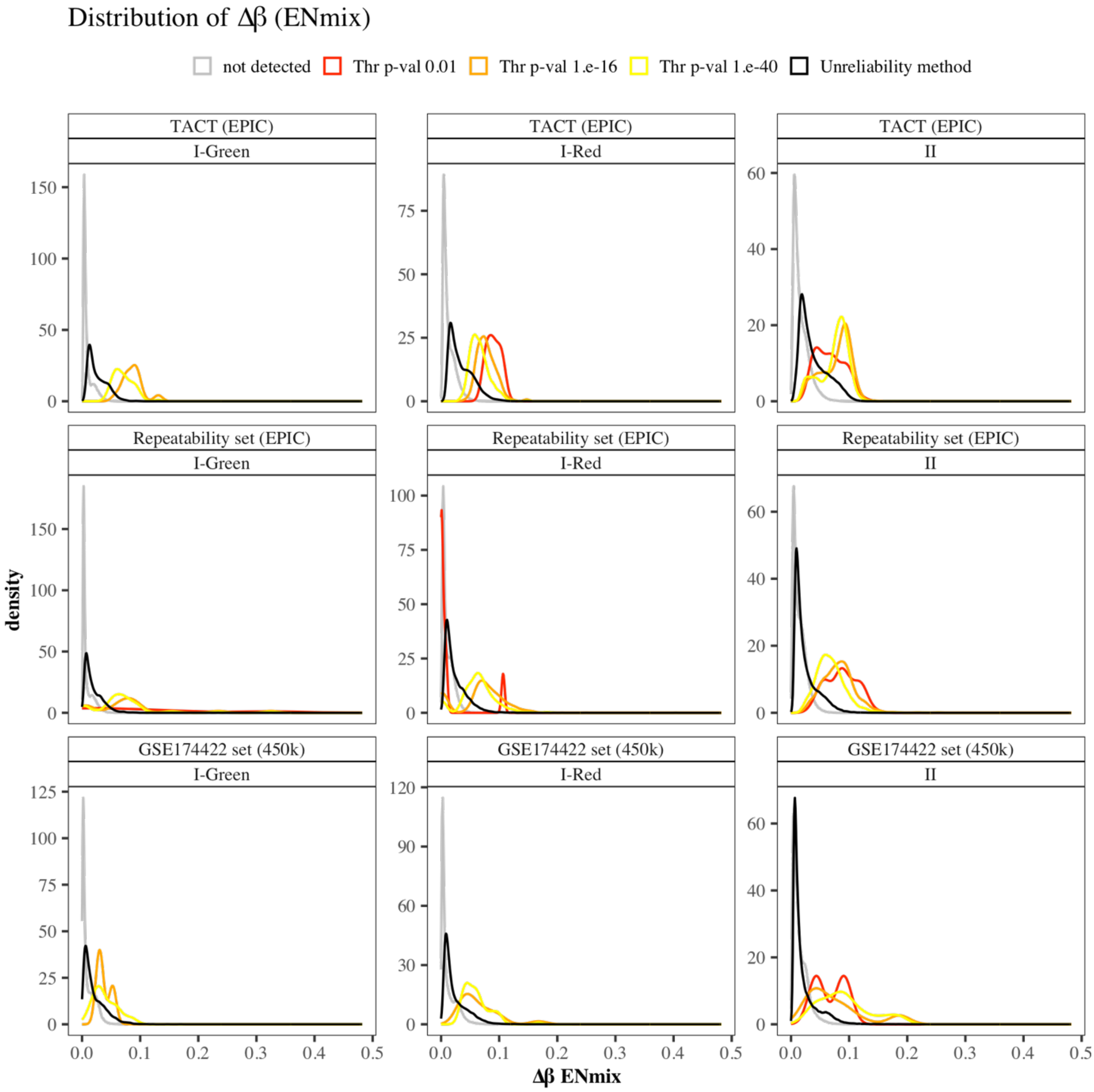


**Figure S10.** Distribution of the averaged, absolute methylation differences in methylation values between repeated samples and technical replicates (Δβ) for all probes (grey) or those removed by the p-value detection method at different threshold settings (red, orange, yellow) and the Unreliability method (black). β-values were normalized using the R package *ENmix*.

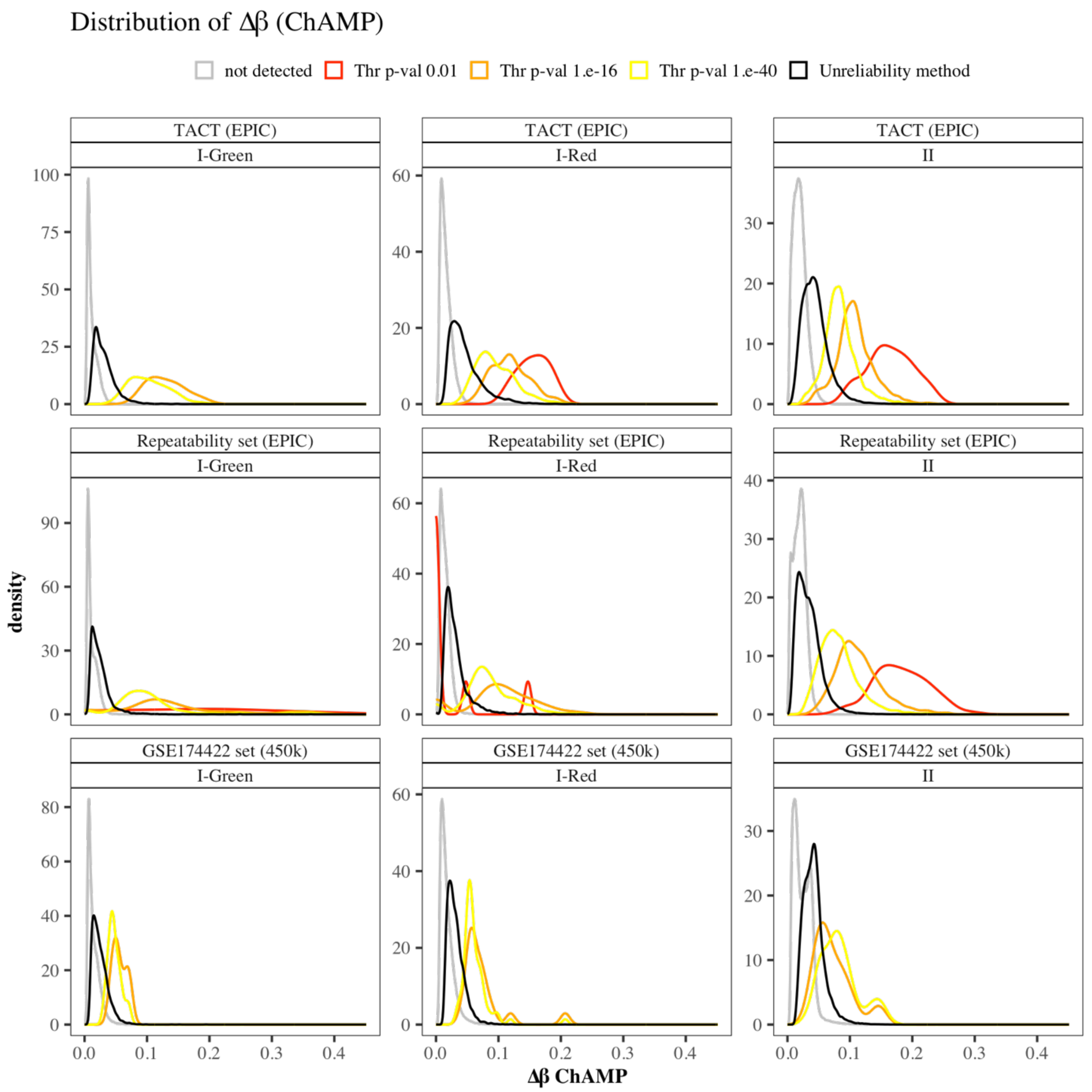


**Figure S11.** Distribution of the averaged, absolute methylation differences in methylation values between repeated samples and technical replicates (Δβ) for all probes (grey) or those removed by the p-value detection method at different threshold settings (red, orange, yellow) and the Unreliability method (black). β-values were normalized using the R package *ChAMP*.

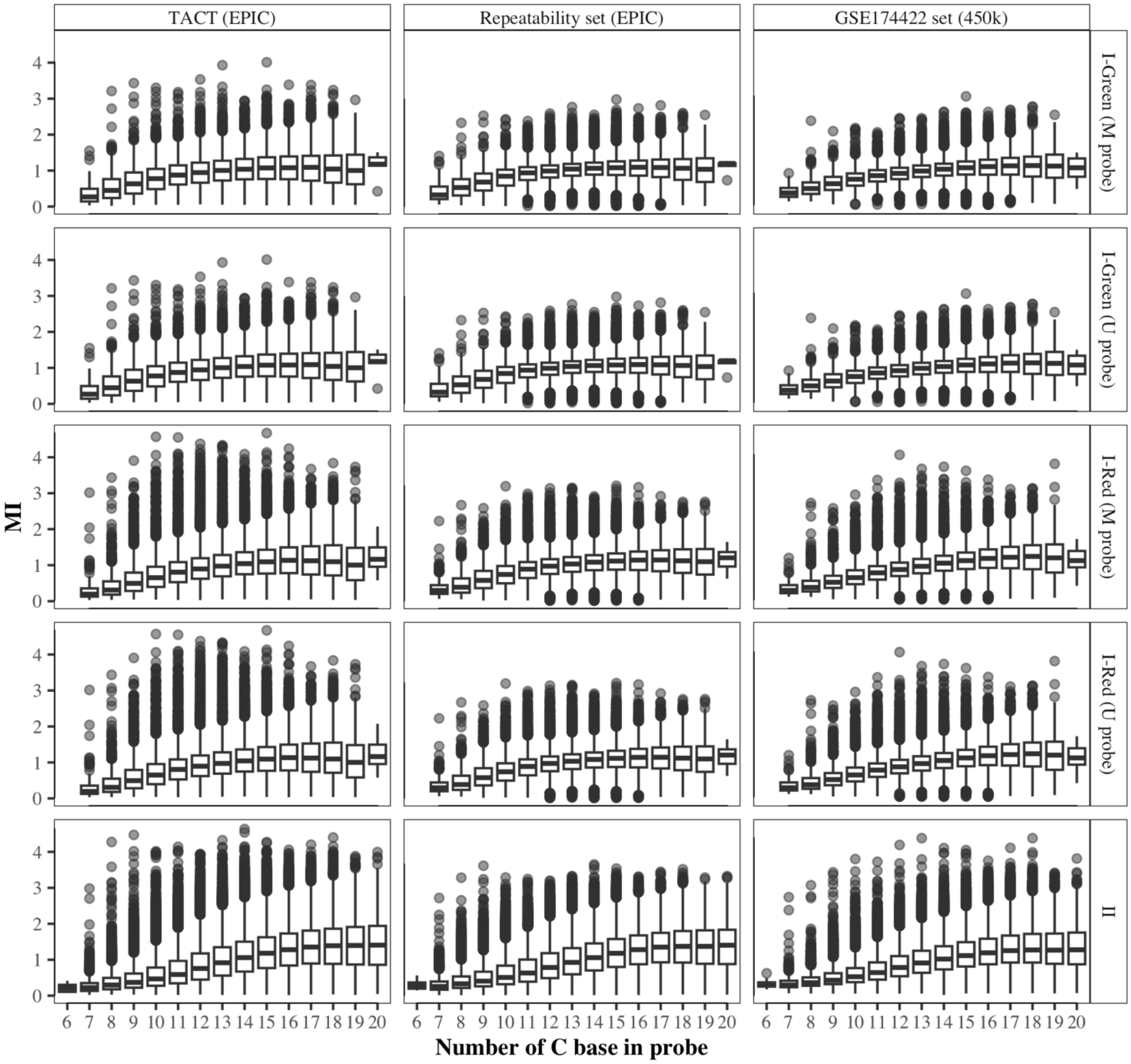


**Figure S12.** Dependence of mean signal intensity (MI) on C content of type I probes in different DNAme data sets.

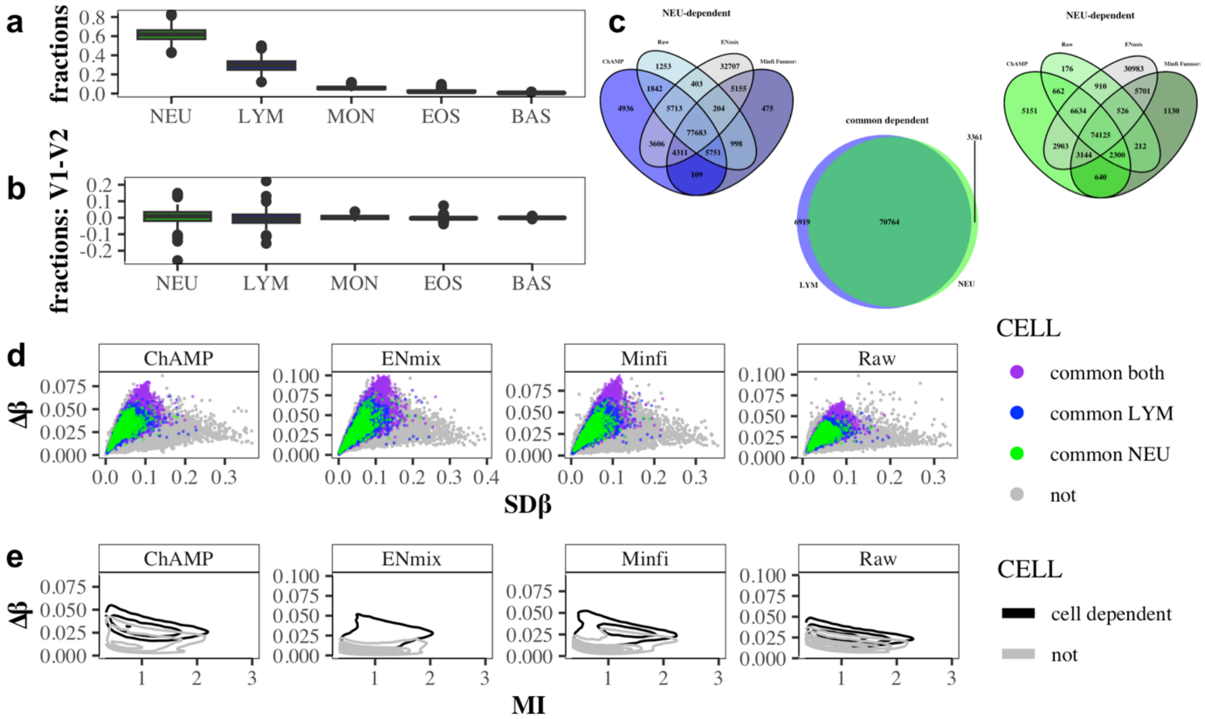

**Figure S13. The effect of cell subtype composition on DNAme variability over time (example for type II probes)(a)** Distribution of cell subtypes fractions in the samples (both visits together); **(b)** Difference in cell subtype fractions for each patient between the two visits; **(c)** Venn Diagram of cell subtype-dependent probes for ChAMP, minfi, and the raw β generation; **(d)** The variability of the probes dependent on one of the cell subtypes fraction, where 'common LYM' – probes significantly detected as lymphocyte-fraction dependent probes for all four pipelines, 'common NEU' – probes significantly detected as neutrophils-fraction dependent probes for all four pipelines, 'common both' – probes significantly detected as neutrophils-fraction dependent and lymphocyte-fraction dependent probes for all four pipelines; **(e)** 2D density plot of variation over time versus MI which shows that cell subtype-dependent probes exhibit high variability in time (which may be interpreted as variability depending on the cellular composition).

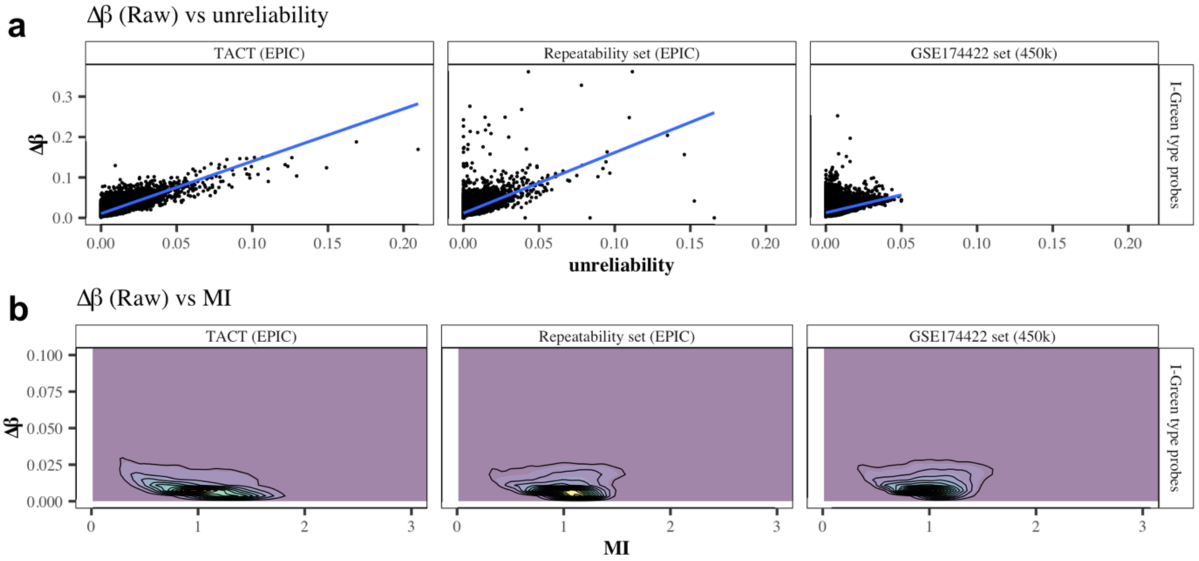


**Figure S14.** Association of type I-Green probe variability (Δβ) with **(a)** the unreliability score and **(b)** the mean intensity (MI) in the TACT study (longitudinal paired blood samples), the Repeatability set (n=4x4x4 technical replicates for fresh blood, frozen blood and cervical smear samples) and GSE174422 (n=2x128 technical replicates for blood samples).

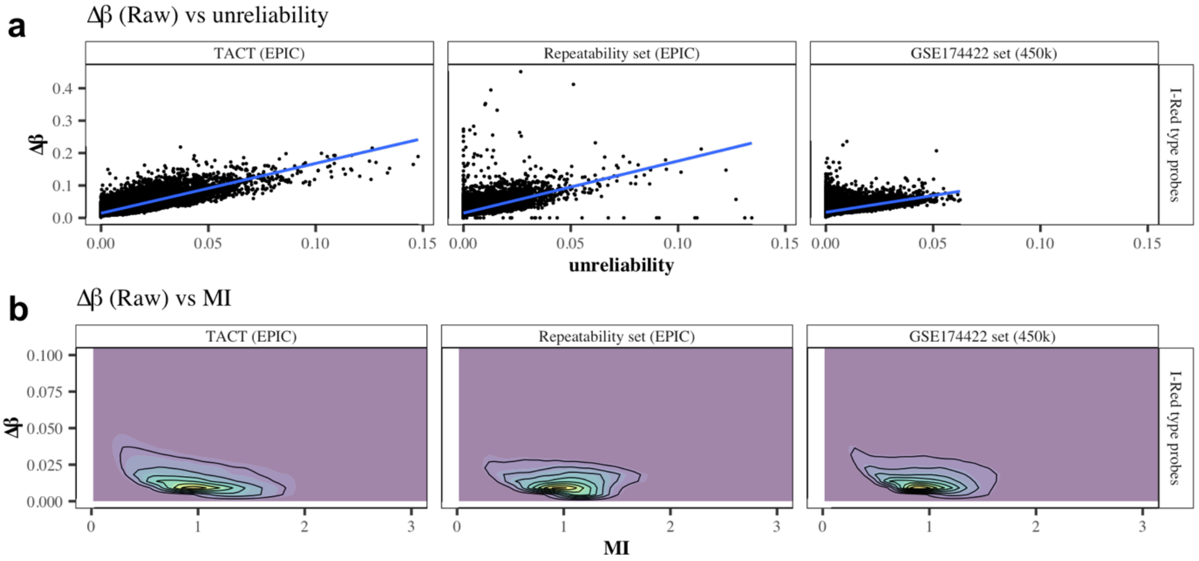


**Figure S15.** Association of type I-Red probe variability (Δβ) with **(a)** the unreliability score and **(b)** the mean intensity (MI) in the TACT study (longitudinal paired blood samples), the Repeatability set (n=4x4x4 technical replicates for fresh blood, frozen blood and cervical smear samples) and GSE174422 (n=2x128 technical replicates for blood samples).

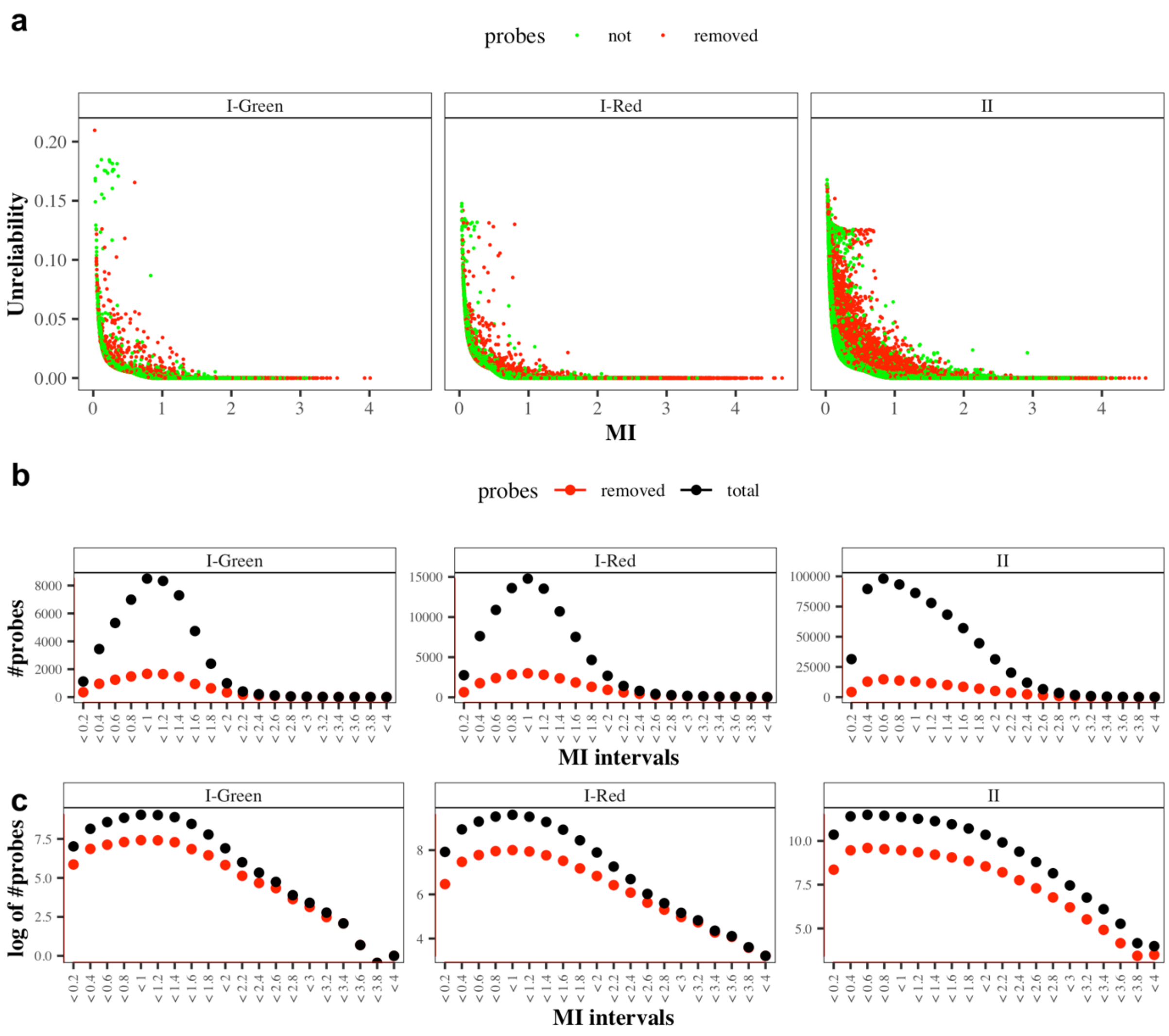


**Figure S15. Probes on MethylationEPIC BeadChip v2.0 retained or removed from v1.0. to with the level of their average normalized intensities (MI). (a)** The removed probes (red dots) do not tend to be low intensity probes and the overall trend for the remaining probes (green dots) still shows tendency to have high unreliability scores on low intensities levels. There is no bias towards the removal of probes with low intensity both in terms of the number of removed probes relative to **(b)** the total number of probes or **(c)** the logarithmic values of the number of probes.

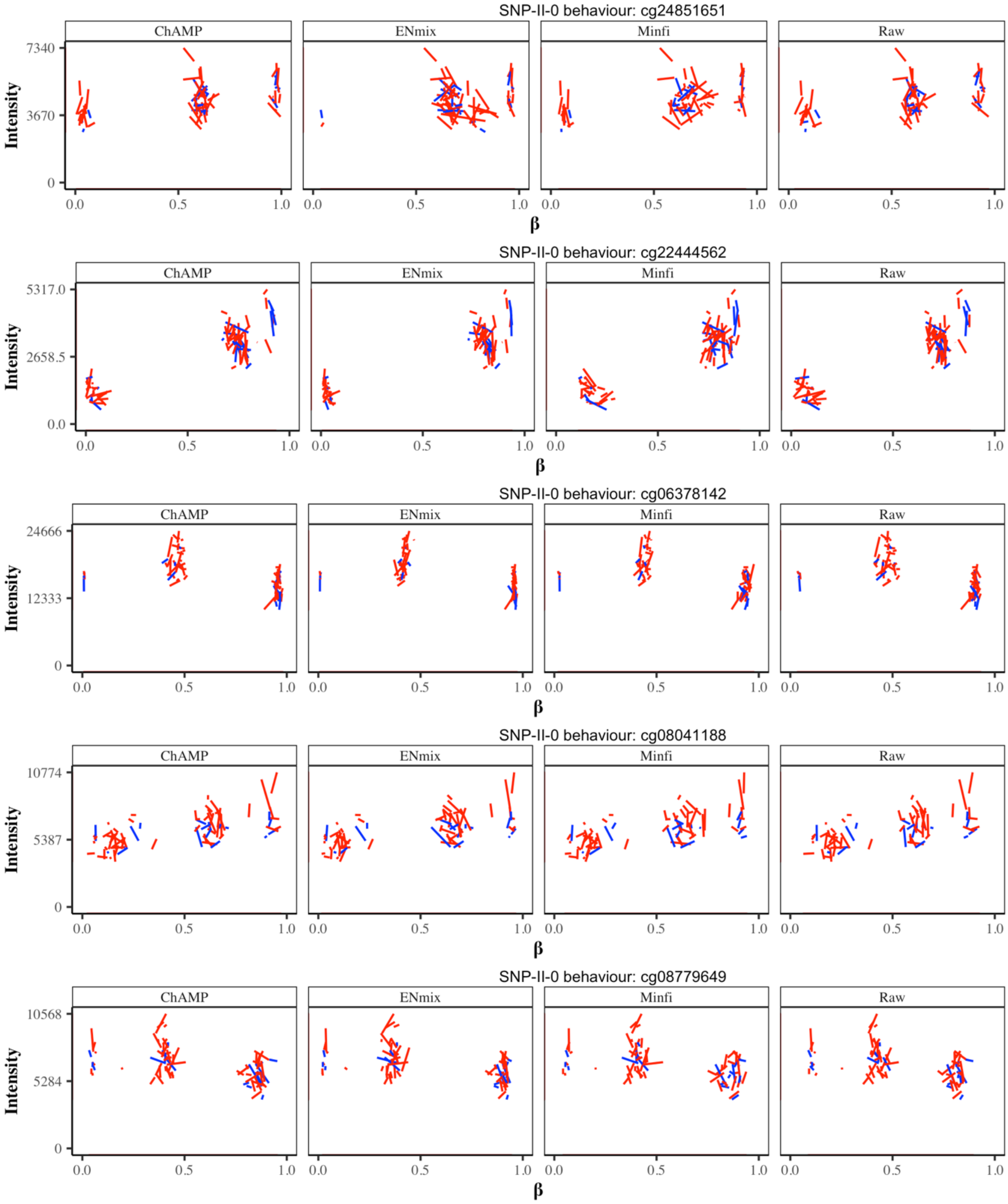


**Figure S16. Examples of type II probes retained on MethylationEPIC BeadChip v2.0 from v1.0 exhibiting SNP-II-0 behavior, but not marked in the manifest.** For clarification on SNP-II-0 behaviour see Figure 1d (main text) and Supplementary Figure S1.

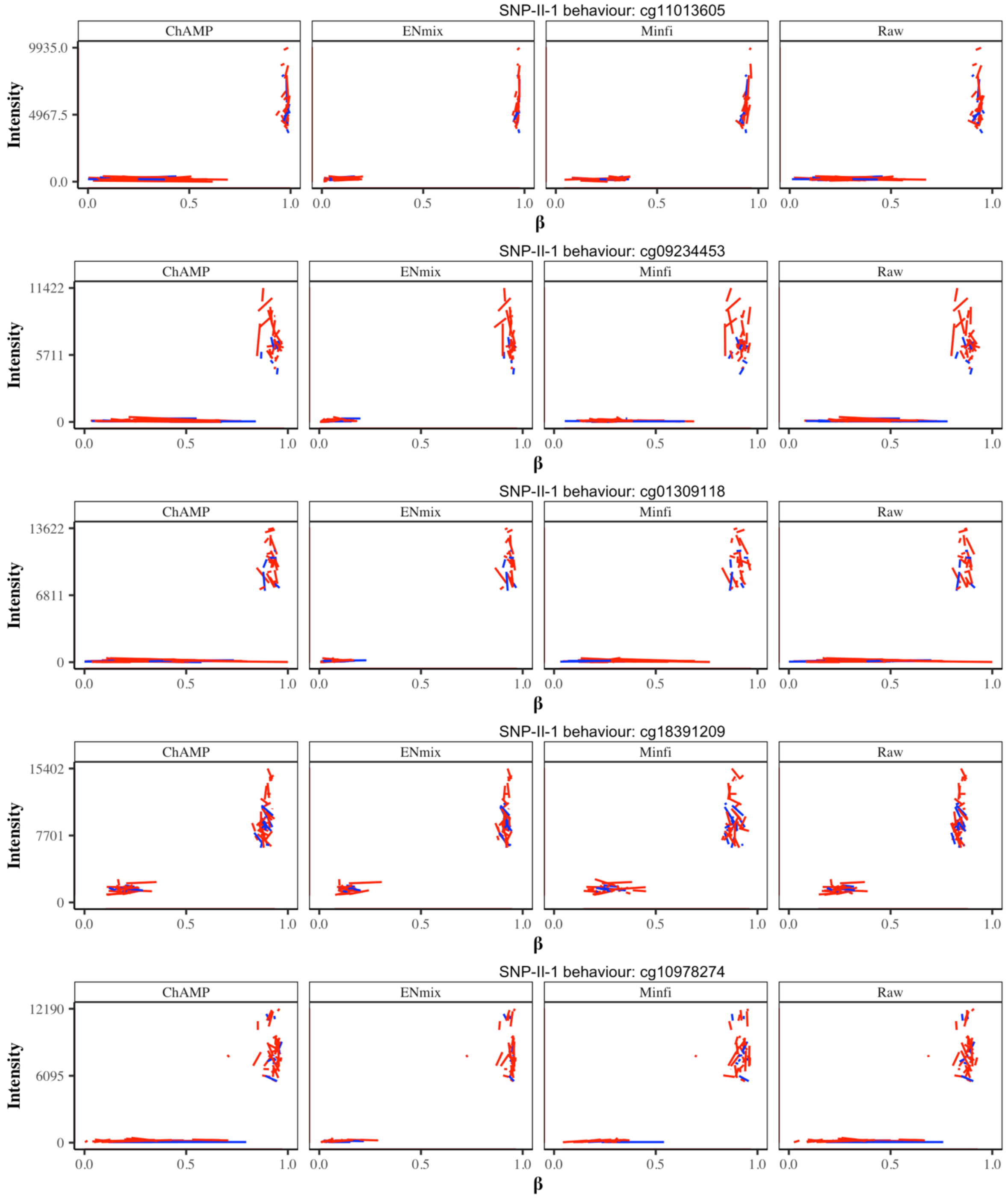


**Figure S17.** ***Examples of type II probes retained on MethylationEPIC BeadChip v2.0 from v1.0 exhibiting SNP-II-1 behavior, but not marked in the manifest.*** *For clarification on SNP-II-1 behaviour see Figure 1d (main text).*
